# Adolescent parvalbumin expression in the left orbitofrontal cortex shapes sociability in female mice

**DOI:** 10.1101/2022.03.04.482801

**Authors:** Yi-Seon Jeon, Daun Jeong, Hanseul Kweon, Jae-Hyun Kim, Choong Yeon Kim, Youngbin Oh, Young-Ho Lee, Chan Hyuk Kim, Sang-Gyu Kim, Jae-Woong Jeong, Eunjoon Kim, Seung-Hee Lee

## Abstract

The adolescent social experience is essential for the maturation of the prefrontal cortex in mammalian species. However, it still needs to be determined which cortical circuits mature with such experience and how it shapes adult social behaviors in a sex-specific manner. Here, we examined social approaching behaviors in male and female mice after post-weaning social isolation (PWSI), which deprives social experience during adolescence. We found that the PWSI, particularly isolation during late adolescence, caused an abnormal increase in social approaches (hypersociability) only in female mice. We further found that the PWSI female mice showed reduced parvalbumin (PV) expression in the left orbitofrontal cortex (OFC_L_). When we measured neural activity in the female OFC_L_, a substantial number of neurons showed higher activity when mice sniffed other mice (social sniffing) than when they sniffed an object (object sniffing). Interestingly, the PWSI significantly reduced both the number of activated neurons and the activity level during social sniffing in female mice. Similarly, the CRISPR/Cas9-mediated knock-down of PV in the OFC_L_ during late adolescence enhanced sociability and reduced the social sniffing-induced activity in adult female mice via decreased excitability of PV^+^ neurons and reduced synaptic inhibition in the OFC_L_. Moreover, optogenetic activation of excitatory neurons or optogenetic inhibition of PV^+^ neurons in the OFC_L_ enhanced sociability in female mice. Our data demonstrate that the adolescent social experience is critical for the maturation of PV^+^ inhibitory circuits in the OFC_L_; this maturation shapes female social behavior via enhancing social representation in the OFC_L_.

**Significance Statement:** Adolescent social isolation often changes adult social behaviors in mammals. Yet, we do not fully understand the sex-specific effects of social isolation and the brain areas and circuits that mediate such changes. Here, we found that adolescent social isolation causes three abnormal phenotypes in female but not male mice: hypersociability, decreased PV^+^ neurons in the OFC_L_, and decreased socially evoked activity in the OFC_L_. Moreover, PV deletion in the OFC_L_ *in vivo* caused the same phenotypes in female mice by increasing excitation compared with inhibition within the OFC_L_. Our data suggest that adolescent social experience is required for PV maturation in the OFC_L_, which is critical for evoking OFC_L_ activity that shapes social behaviors in female mice.

## Introduction

One of the hallmarks of postnatal maturation in the mammalian cortex is the maturation of GABAergic inhibitory circuits (Luhmann and Prince, 1991). Maturation is accompanied by an increase in the expression of both the parvalbumin (PV) in a subset of GABAergic neurons and the perineuronal nets (PNNs) surrounding the PV^+^ neurons (Celio and Blumcke, 1994; Huang et al., 1999). PV is a calcium-binding protein and a well-known genetic marker for fast-spiking GABAergic neurons in the cortex (Rudy et al., 2011). PV gene expression is upregulated in the sensory cortex in an activity-dependent manner (Sugiyama et al., 2008; Krishnan et al., 2017) after mammals are born. Like PV, PNNs also increase during cortical maturation and, in the adult cortex, surround most of the PV^+^ neurons (Ueno et al., 2017; Ueno et al., 2018). Along with the increase of PNNs in cortical circuits, other molecular changes reduce net plasticity and facilitate the maturation of the cortical network (Beurdeley et al., 2012). If mice are deprived of visual input during the critical period after eye opening, via monocular deprivation or dark rearing, PV^+^ neurons are less developed, and PNNs are reduced in the primary visual cortex (V1) (Sugiyama et al., 2008). This deprivation, in turn, diminishes both the visual response of V1 neurons that receive input from the deprived eye (Sugiyama et al., 2008) and inhibition in the cortex (Dityatev et al., 2007). Therefore, the postnatal sensory experience is critical for the expression of PV and PNNs in PV^+^ neurons, as these are required for the maturation of PV^+^ inhibitory circuits and, eventually, for shaping sensory responses in the cortex (Hensch, 2005).

The maturation of the cortex occurs first in the sensory area, where the process is part of sensory experience, and later in the motor-frontal and the PFC (Hensch, 2005). The social experience is critical for the maturation of the PFC in both humans (Blakemore, 2008; Paus et al., 2008) and rodents (Makinodan et al., 2012; Bicks et al., 2020; Tan et al., 2021), particularly during juvenile and adolescent periods. Patients with autism or schizophrenia often show abnormal social cognition as well as dysfunction in the PFC (Bicks et al., 2015). In these patients, the expression of PV and PNNs is reduced in the dorsolateral PFC (Enwright et al., 2016). Thus, it is plausible that immature PV^+^ circuits in the PFC cause the psychiatric symptoms underlying abnormal social behavior in mammals. However, we still do not understand whether social experience is required for PV^+^ inhibitory circuits in these PFC regions to mature. In particular, it is largely unknown whether social experience affects the maturation of inhibitory circuits in the orbitofrontal cortex (OFC), one of the PFC sub-regions critical for social cognition (Adolphs, 2003; Amodio and Frith, 2006).

Mammalian social behavior is important for maintaining a conspecific society. Social interaction during adolescence is critical for shaping social and cognitive behaviors in both humans and rodents (Spear, 2000; Makinodan et al., 2012; Hinton et al., 2019; Almeida et al., 2021; Potrebic et al., 2022). Interestingly, the mammalian brain shows hemispheric asymmetry and lateralization in executing social behaviors (Marlin et al., 2015). Even in rodents, according to recent studies, the PFC functions asymmetrically in mediating social behaviors (Lee et al., 2015; Marlin et al., 2015). However, whether such asymmetry in the PFC becomes increasingly pronounced in a sex-specific manner with social experience during adolescence is still unclear. We investigated the effect of adolescent social isolation on the maturation of PV^+^ neurons in the PFC in both sexes and hemispheres. Using post-weaning social isolation (PWSI) and the CRISPR/Cas9 system, we revealed that social isolation during adolescence caused the defect in PV expression in the OFC_L_ of female mice, which led to hypersociability and reduced OFC_L_ activity during social contact.

## Materials and Methods

### Experimental model and subject details

We approved all experimental procedures from the KAIST Institutional Animal Care and Use Committee (IACUC-18-236). We maintained all mice ad libitum under light from 8 am to 8 pm and dark cycle from 8 pm to 8 am. All mice were weaned at postnatal day 21 (P21); mice were reared in double (28cm × 17cm × 14cm) cages with 3-5 mates for group housing or in single (20cm × 10cm × 14cm) cages alone for social isolation. We performed experiments on WT (C57BL/6J), PV-Cre (JAX, #008069), and the Rosa26-floxed STOP-Cas9 knock-in (JAX # 024857) mice.

### Histology

We anesthetized mice with isoflurane and then perfused them with 1X phosphate-buffered saline (PBS) and 4% paraformaldehyde (PFA, w/v in PBS). Mouse brain samples were post-fixed for 4h at 4°C; after fixation, we incubated them in 30% sucrose solution (w/v in PBS) for 1-2 days at 4°C. Then, we embedded each brain sample in an optimal cutting temperature medium (OCT, Tissue-Tek, Cat#4583) and froze each at −80°C. We sectioned frozen brains using a cryostat (Leica) and collected coronal sections either at 30 μm (for confirming injection sites) or 15 μm (for immunostaining) thickness. To confirm fluorescently labeled injection sites, brain sections were washed three times with 1X PBS and mounted in DAPI-containing mounting medium (VECTASHIELD®, cat#H-1200). For immunostaining, we washed the brain sections three times with 1X PBS and permeabilized them with 0.3% Triton X-100 in 1X PBS for 30 min. After the sections were incubated in a blocking solution (2% normal donkey serum in 1X PBS) for 2 hours at room temperature, we incubated them with primary antibodies (rabbit anti-PV, 1:500 dilution in blocking solution, Swant, cat#PV25; mouse anti-NeuN, 1:500 dilution in blocking solution, Millipore, cat#MAB377) for 2 days at 4 °C. Last, we incubated them with fluorescence-tagged secondary antibodies (Alexa Fluor 594 donkey anti-rabbit IgG for PV staining, 1:500 dilution in 1X PBS, Life Technology, cat#A21207; Alexa Fluor 647 donkey anti-mouse IgG for NeuN staining, 1:500 dilution in 1X PBS, Jackson Immunoresearch, cat#715-605-151). For PNN staining, we treated the brain sections with the Fluorescein (FITC) labeled Wisteria Floribunda Lectin (WFA, WFL) (1:500 dilution in 1X PBS, Vector labs, cat#FL-1351) for 2 hours at RT before mounting them in clear mounting medium (DAKO, cat#S3023). We imaged fluorescent signals of mounted brain sections under a scanning microscope (ZEISS, Axio Scan.Z1).

### Cell counting analysis

When we sectioned the mouse brain samples, we collected 15μm brain sections in every 60μm thickness across the frontal cortex. After immunostaining the sections, we scanned the fluorescent images of the brain sections and matched the images to the mouse brain atlas (Franklin and Paxinos) to confirm the anterior-to-posterior location of the coronal brain sections. We then selected three images of sections in a row that include OFC areas (between +2.10mm ~ +2.34mm anterior from the bregma) (see Fig 2C). To count the labeled neurons in the OFC, we put a 0.3 × 0.3mm^2^ square as a region of interest (ROI) at the center of the OFC (±1.3mm lateral from the bregma; depth, −2.5mm). To quantify the number of PV^+^ and NeuN^+^ neurons, we used the automatic cell counting software (ImageJ) (Grishagin, 2015). After converting the fluorescent images to the 8-bit grayscale images, we further transformed the grayscale images into binary images. For the binary transition, we used distinct thresholds at the grayscale for isolating the NeuN^+^ neurons (Fig. 2, 11.39±0.02% of total mean intensity) and the PV^+^ neurons (Fig. 2, 2.12±0.16% of total mean intensity). We then automatically counted the white-filled circles as cells if they satisfied the following conditions: 1) the total size is above 60 × 60 pixels, and 2) the circularity is above 0.2. We set the thresholds in each sample at the level that yielded a similar number of cells from the automatic counting compared to the manual counting using a couple of template images from each sample. There was no significant difference between the number of cells counted automatically and the number of cells counted manually in a blind manner (see Fig. 2B). To validate the knock-down efficiency of sgPV or chABC (see Fig 5), we manually and blindly counted the number of PV^+^ or WFA^+^ neurons in the ROI.

### Sociability test

We performed the sociability test by using two types of mazes: one is a modified 3-chamber-like maze (50cm × 45cm × 20cm) where two containers were located diagonally at opposite corners, and the other is a standard 3-chamber maze (60cm × 40cm × 20cm), which was used in other literature (Won et al., 2012). Most of the data we plotted in the manuscript were collected in the modified maze. We performed the sociability test on mice at either P79-81 or P50. Before behavior experiments, all mice were habituated for 30-60 minutes in the habituation chamber. All mice were maintained in a group-housed or an isolated condition until the habituation. Mice were then moved to the sociability test maze. The sociability test consisted of two phases: 1) habituation in the sociability maze (empty containers, 10min) and 2) sociability test (an object container and a social container with a stranger mouse, 10min). We used a 2-week younger and same-sex mouse as a stranger mouse not to evoke any aggressively or sexually driven social behaviors in the tested mouse (Murugan et al., 2017). We used the software, EthoVision®XT 11.5 (Noldus), to automatically analyze interaction times and manually analyze the sniffing time. The sniffing time was defined as the time when the subject mouse’s nose contacted the container either empty or with a novel mouse. We analyzed mouse sociability by two indicators: time spent in interaction zones near the container (18cm × 18cm) and sniffing time. We calculated the social sniffing and social interaction indices as follows (Nakanishi et al., 2017; Cheong et al., 2020):

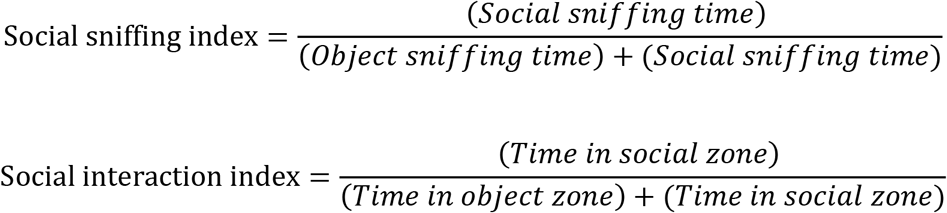

### Resident-intruder (RI) test

We devised the resident-intruder (RI) test as previously described (Tan et al., 2021). The mouse was single-housed for 24hr and then exposed to an intruder mouse in the home cage for 10 minutes. We used a slightly smaller (10-20% lighter) unfamiliar sex- and strain-matched novel mouse as an intruder. The offensive behaviors of the subject mouse against the intruder, including lateral threat, upright posture, clinch attack, and chase, were scored as aggressive behaviors. In addition, the non-offensive social behaviors in the RI test, such as nose-to-nose interaction and anogenital sniffing, were scored. All scoring was performed blindly.

### Measuring estrous cycle in female mice

We monitored the estrous cycle of female mice using vaginal cytology (McLean et al., 2012). Vaginal smears from subject mice were collected right after the sociability test. Smears were placed on slides and stained with 0.1% crystal violet (Sigma Aldrich, cat#V5265). Each estrous cycle stage was determined based on the cell type that was mainly observed in the smear sample among the three types: nucleated epithelial cells, cornified squamous epithelial cells, and leukocytes. To examine the difference in estrous cycles between GH and PWSI, we performed the vaginal cytology for four consecutive days in each mouse.

### *In vivo* extracellular recordings for measuring sniffing-induced activity

We performed all recordings in the head-fixed and awake mice. We marked the top of left and right OFCs and implanted custom-designed head-plates on the skull of mice using dental cement and small screws. After 2-3 days of recovery, we habituated mice to be head-fixed for 20min per day for 3 days. After the habituation, we performed the recording in the OFC by making a small craniotomy on the marked spots and inserting 32 channel silicon probes (A1X32-Poly2-10mm-50-177 or A1X32-Poly3-10mm-50-177, Neuronexus) vertically into each OFC using a micro-drive manipulator (Siskiyou). We first recorded spiking activity in the OFC without presenting a stranger mouse for 10 min (object_sniffing_ session) and next continuously recorded neural activity while presenting a head-fixed stranger mouse in front of the recorded mouse for 10 min (social_sniffing_ session). During the recording, we monitored and recorded the sniffing behavior of the mouse by using a web camera (Microsoft, LifeCam HD-3000). All extracellular signals were recorded at a 30kHz sampling rate, filtered between 500~5000Hz, and amplified by a miniature digital headstage (CerePlex μ, Blackrock Microsystems). After the recording, we confirmed the recording sites by injecting red retrobeads (Lumaflour) into the recording areas and performing histology. If the recording sites were outside of the OFC, we did not use those recording data in further analysis.

#### Automatic quantification of sniffing behaviors

We developed a Matlab code to automatically analyze sniffing events during object- and social sniffing sessions in head-fixed mice. First, we converted all the RGB image frames of a recorded video clip to grayscale using the *‘rgb2gray’* function. Next, we chose the region of interest (ROI), including the nose and whiskers of the mouse being recorded, made a binary matrix with 1 inside the ROI and 0 outside the ROI of each pixel, and multiplied an original image by the binary matrix of each corresponding pixel along all the frames (ROI_1). We subtracted background brightness from ROI_1 to eliminate overall brightness fluctuations (ROI_2) and calculated deviations of brightness within the ROI by subtracting the mean brightness of each pixel (ROI ΔF). To detect sniffing events, we subtracted a moving average (F_0_, sliding window = 50 frames) from ROI ΔF (ROI ΔF–F_0_), and set a threshold by multiplying a median of absolute ROI ΔF–F_0_ by 4. If ROI ΔF–F_0_ in each frame crossed the threshold, we included the frame in a sniffing event.

#### Neural activity analysis

We performed the spike-sorting using Klusters software (http://neurosuite.sourceforge.net/) (Hazan et al., 2006). We grouped 32-channel data into eight groups and sorted single units by principal components analysis. We only isolated single units that show more than a 2 ms refractory period in their auto-correlogram. We performed all other analyses using custom-made codes in Matlab (Mathworks). We sorted sniffing trials in object and social sessions by identifying sniffing events across the recording sessions. We only used trials that satisfied the following two criteria: 1) inter-sniffing interval (time between the offset of sniffing in the current trial and the onset time of the next trial) > 5sec, and 2) sniffing duration (time between the onset and the offset of sniffing in one trial) > 1sec. To plot the peri-event time histogram (PETH) of all neurons as a population, we performed Z-scoring by averaging firing rates (Hz) of individual units across trials and normalized them by 1sec baseline activity (3 to 2 sec before the onset of sniffing). The normalized firing rates were convolved with a Gaussian filter (σ = 50ms) only for visualization purposes. To identify the neurons showing significant sniffing-induced activity, we statistically compared the firing rates of individual neurons during the 2-sec pre-period (3 to 1 sec before the onset of sniffing) and 2-sec post-period (0 to 2 sec after the onset of sniffing) of each sniffing trial by bootstrapping (p < 0.01, n = 5000). We quantified the sniffing-induced activity of neurons by averaging Z values from 0 to 2 sec after the onset of sniffing. We classified cell types of recorded neurons based on spike waveforms by quantifying spike widths of each isolated neuron (time between peak and through of the spike, μs). If the spike width was below 350 μs or above 450 μs, recorded neurons were classified as fast-spiking (FS) neurons or regular-spiking (RS) neurons, respectively. For the baseline activity analysis, we quantified the average firing rates (Hz) from 0 to 2 sec before the onset of sniffing.

### Perineuronal net degradation

To remove the PNN, we reconstituted protease-free chABC (Amsbio) in PBS to a final concentration of 61U/ml. We anesthetized mice by inhalation of 1.5 % isoflurane in oxygen on the stereotaxic apparatus. We injected 0.5μl of PBS or chABC into the left OFC (AP+2.35, ML+1.3, DV −2.5) of female mice at postnatal day 70-75. We performed behavior experiments seven days after the injection and sacrificed the mice to confirm PNN degradation by histology. To stain the PNN in the brain slice containing the OFC_L_, we treated the fluorescein (FITC)-conjugated Wisteria Floribunda Lectin (WFA, WFL) (Vector labs, cat#FL-1351) as a secondary antibody, diluted at 1:500 in 1X PBS, for 2 hours at RT before mounting the slices in a transparent mounting medium (DAKO, cat#S3023). We manually and blindly counted the number of WFA^+^ neurons within 0.5 × 0.5mm^2^ ROIs in three representative coronal sections covering OFC.

### CRISPR / Cas9 experiments

#### Generating the PX552-sgPV-tdTomato

To generate the AAV-tdTomato vector, we first replaced the eGFP gene in PX552 (Addgene # 60958) with tdTomato. The tdTomato template was obtained from pAAV-FLEX-tdTomato (Addgene # 28306). Next, we cloned two AAV-tdTomato plasmids, both of which induce the targeted mutations at the parvalbumin gene (sgPV). One virus expresses two sgRNAs targeting parvalbumin exon4: sgRNA1, 5’-AGACAAGTCTCTGGCATCTG-3’ and sgRNA2, 5’-AAGGATGGGGACGGCAAGAT-3’ (Cloned recombinant DNA fragment: AGACAAGTCTCTGGCATCTGGTTTTAGAGCTAGAAATAGCAAGTTAAAATAAGGCTAGTCCGTTATCAACTTGAAAAAGTGGCACCGAGTCGGTGCAACAAAGCACCAGTGGTCTAGTGGTAGAATAGTACCCTGCCACGGTACAGACCCGGGTTCGATTCCCGGCTGGTGCAAAGGATGGGGACGGCAAGATGTTTTAGAGCTAGAAATAGCAAGTTAAAATAAGGCTAGTCCGTTATCAACTTGAAAAAGTGGCACCGAGTCGGTGC, Bioneer & In-house PCR amplification), and the other virus expresses two sgRNAs targeting parvalbumin exon5: sgRNA3, 5’-GCTAAGTGGCGCTGACTGCT-3’ and sgRNA4, 5’-GGCCGCGAGAAGGACTGAGA-3’(Cloned recombinant DNA fragment: GCTAAGTGGCGCTGACTGCTGTTTTAGAGCTAGAAATAGCAAGTTAAAATAAGGCTAGTCCGTTATCAACTTGAAAAAGTGGCACCGAGTCGGTGCAACAAAGCACCAGTGGTCTAGTGGTAGAATAGTACCCTGCCACGGTACAGACCCGGGTTCGATTCCCGGCTGGTGCAGGCCGCGAGAAGGACTGAGAGTTTTAGAGCTAGAAATAGCAAGTTAAAATAAGGCTAGTCCGTTATCAACTTGAAAAAGTGGCACCGAGTCGGTGC, Bioneer & In-house PCR amplification).

After cloning the AAV-tdTomato vectors, we transfected them with packaging plasmid into HEK293T (KCTC, cat# HC20005) to produce AAV-sgPV-tdTomato (exon4 and exon5) and AAV-tdTomato, using lipofectamine 2000 (Invitrogen, cat# 11668-019). After 72hr, we collected AAV (AAV2) particles from the cell lysates and performed iodixanol gradient ultracentrifugation. Next, we concentrated the viral particles with Amicon Ultra Centrifugal Filter (Millipore) and quantified the concentration of viral particles by quantitative PCR (genome copies per ml; AAV-tdTomato, 3.92 × 10^14^; AAV-sgPV(exon4)-tdTomato, 3.07 × 10^12^; AAV-sgPV(exon5)-tdTomato, 7.17 × 10^12^).

#### Validation of CRISPR/Cas9 in the Neuro2a cell

To identify the efficiency of sgPV in knocking down the PV, we transfected cloned AAV vectors with the Cas9-eGFP expressing vector (PX458) in the Neuro2a cells. After 72h, we collected and sorted tdTomato/eGFP double-positive cells using BD FACSAria II (BD Biosciences). Next, we purified genomic DNA from sorted cells and performed PCR to amplify the target sites of the PV gene, adding the index sequence for sequencing. Last, we subjected the samples to Miniseq (Illumina) and analyzed insertion and deletion frequencies using CRISPR RGEN Tools (http://www.rgenome.net/).

#### Behavior and physiological experiments with CRISPR/Cas9 mice

To express Cas9 selectively in PV^+^ neurons, we crossed the PV-Cre mice (JAX #008069) with the Rosa26-floxed STOP-Cas9 knock-in mice (JAX # 024857). We injected AAV-tdTomato (tdTom) or AAV-sgPV-tdTomato (sgPV) into the crossed mice (PV::Cas9) on the postnatal day 50-65. On postnatal days 78-85, we performed the behavior test and *in vivo* recording. To verify whether the AAV expression knocked down the PV effectively, we collected brain samples of the injected mice and performed immunohistochemistry as described above. We used a combination of the primary (rabbit anti-PV for PV staining, goat anti-GFP for Cas9 staining) and the secondary (Alexa Fluor 405 donkey anti-goat IgG and Alexa Fluor 647 donkey anti-rabbit IgG were used) antibodies.

#### Whole-cell patch-clamp recordings in brain slices

For electrophysiology experiments, tdTom/sgPV-expressed brain samples were collected from anesthetized PV::Cas9 mice with isoflurane (Terrell) and sliced (coronal, 300 μm thickness) using a vibratome (Leica VT1200). Brain extraction and sectioning were conducted in ice-cold dissection buffer containing (in mM) 212 sucrose, 25 NaHCO_3_, 5 KCl, 1.25 NaH_2_PO_4_, 0.5 CaCl_2_, 3.5 MgSO_4_, 10 D-glucose, 1.25 L-ascorbic acid and 2 Na-pyruvate bubbled with 95% O_2_/5% CO_2_. The sectioned slices were incubated in a recovery chamber at 32 °C with normal ACSF (in mM: 125 NaCl, 2.5 KCl, 1.25 NaH_2_PO_4_, 25 NaHCO_3_, 10 glucose, 2.5 CaCl_2_ and 1.3 MgCl_2_, oxygenated with 95% O_2_/5% CO_2_). After 60-min recovery at 32 °C, slices were incubated for 30 min at room temperature (20–25 °C). The slice was transferred to a recording chamber at 27–29 °C with circulating ACSF saturated with 95% O_2_ and 5% CO_2_. Recording pipettes were made from thin-walled borosilicate capillaries (30–0065, Harvard Apparatus) with resistance 3.0–4.0 MΩ using a two-step vertical puller (PC-10, Narishege).

Whole-cell patch-clamp recordings were conducted using a MultiClamp 700B amplifier (Molecular Devices) and Digidata 1550 (Molecular Devices). We recorded both AAV-expressed and non-expressed neurons in the OFC_L_. We confirmed the AAV expression by imaging the red fluorescence of the patched cells under the fluorescent microscope (Olympus). During the recordings, hyperpolarizing step pulses (5mV, 40ms) were injected into the cell to monitor membrane resistance calculated by the peak amplitude of the capacitance current. Data from cells that changed the resistance by more than 20% were excluded. To measure the intrinsic excitability, recording pipettes (3.0–4.0MΩ) were filled with an internal solution containing (in mM) 137 K-gluconate, 5 KCl, 10 HEPES, 0.2 EGTA, 10 Na-phosphocreatine, 4 Mg-ATP, and 0.5 Na-GTP, with pH 7.2, 280 mOsm. Picrotoxin (Sigma, 100μM), NBQX (Tocris, 10μM), and D-AP5 (Tocris, 50μM) were added to ACSF to inhibit postsynaptic responses. After gigaseal and rupture, currents were clamped, and RMP was maintained at −70mV. To measure the I-V curve, current inputs were increased from −300 to 120pA in increments of 30pA per sweep with a time interval of 2 seconds. To measure the I-F curve, current inputs were increased from 0 to 660pA in increments of 60pA per sweep with a time interval of 10 seconds. For measuring sEPSCs in the OFC PV^+^ neurons, glass pipettes were filled with an internal solution composed of (in mM) 100 CsMeSO_4_, 10 TEA-Cl, 8 NaCl, 10 HEPES, 5 QX-314-Cl, 2 Mg-ATP, 0.3 Na-GTP, and 10 EGTA, with pH 7.25, 295 mOsm at a holding potential of −70 mV. Picrotoxin (100μM) was added to ACSF to block IPSCs. For measuring sIPSCs in the OFC excitatory neurons, recording pipettes were filled with an internal solution containing (in mM) 120 CsCl, 10 TEA-Cl, 8 NaCl, 10 HEPES, 5 QX-314-Cl, 4 Mg-ATP, 0.3 Na-GTP, and 10 EGTA, with pH 7.35, 280 mOsm holding at −70 mV. Circulating ACSF were added NBQX and D-AP5 to block EPSCs. Data were acquired by Clampex 10.2 (Molecular Devices) and analyzed by Clampfit 10 (Molecular Devices).

### Wireless optogenetic modulation of the OFC neurons *in vivo*

#### Design and fabrication of flexible optoelectronic neural probes

Flexible optoelectronic probes were designed using electronic design automation software (Altium Designer 18.0, Altium Limited) and produced using a conventional photolithography process. Neural probes (0.1 mm thick) were designed in a bilateral form, in which the right and the left probes (0.38 mm wide and 4 mm long each) were separated by 2.6 mm, to access both regions of the mouse brain. Copper traces and electrodes (18 μm-thick) were patterned on one side (i.e., top layer) of the polyimide substrate (25 μm-thick). In contrast, the other side (i.e., the bottom layer) was covered with a copper layer (18 μm-thick) to stiffen the probe for penetration into brain tissue. The probe tip part, where the microscale inorganic light-emitting diodes (μ-ILEDs; TR2227, Cree; blue (473 nm) or green (532 nm), 270 × 220 × 50 μm^3^) were attached, was not covered with a copper layer to allow light emission from both sides of the probe. The fabricated probes showed an optical intensity of ~405 mW/mm^2^ at the top layer and ~45.3 mW/mm^2^ at the bottom layer, which sufficiently exceeds the threshold intensity for ChR2 activation (~1 mW/mm^2^). A 3-pin plug-n-play female pin connector (M50-3130345, Harwin) and two μ-ILEDs were mounted on the top layer electrodes after applying a low-temperature solder paste (SMDLTLFP10T5, Chip Quik Inc.) and soldered using a precision soldering iron (NASE-2C, JBC Soldering Tools) with the temperature of 250 °C. After confirming that every component was robustly attached, probes were coated with Parylene C (7 μm-thick) for waterproofing. Lastly, the probes were vertically bent and attached to a 3D-printed probe holder, followed by fixing them in place using epoxy (5 Minute Epoxy, Permatex).

#### Design of programmable wireless control module with detachable configuration

A Bluetooth Low Energy (BLE) based wireless control module was designed to be easily attached and detached with a bilateral optoelectronic probe and a battery through a plug-n-play connection ((Qazi et al., 2019; Lee et al., 2020). This module controls the optical output of the μ-ILEDs through smartphone manipulation. The electronic circuit of the wireless module consists of a rechargeable Lithium Polymer (LiPo) battery (GM300910-PCB, PowerStream Technology; 12 mAh, 9 × 10 × 3 mm^3^), a BLE System-on-Chip (SoC; EYSHSNZWZ, Taiyo Yuden; 8.55 × 3.25 × 0.3 mm^3^), and various other electronic components (a linear voltage regulator, four decoupling capacitors, two resistors, and two indicator LEDs). The current supplied from the LiPo battery is regulated and stabilized by a voltage regulator (NCP4624DMU30TCG, onsemi) and decoupling capacitors and flows to the BLE SoC to operate the module. Two indicator LEDs (VAOL-S4RP4, VCC; red (624 nm wavelength)) were placed at the output pins of BLE SoC, each connected in series with a resistor (RK73H1HTTC6041F, KOA Speer; 6.04 kΩ), to visually show the current operation status of the module during the animal experiment – for example, the indicator LED blinks every 5 seconds during optical stimulation. The wireless control module can be assembled with optoelectronic probes through 3-pin plug-n-play connectors (male: M50-3630342, Harwin; female: M50-3130345, Harwin) and assembled with LiPo battery through 2-pin connectors (male: M50-3630242, Harwin; female: M50-3130245, Harwin). Every component except the LiPo battery was soldered on the wireless module flexible printed circuit board (FPCB) of the wireless module (9.3 × 7.6 × 0.1 mm^3^), which was designed and manufactured through the same method for the μ-ILED probes. The module was wirelessly programmed over the air (OTA; Nordic Device Firmware Update (DFU) bootloader) to provide a pulse (20 Hz with 25 ms pulse width; i.e., 50% duty cycle) or continuous (i.e., 100% duty cycle) current to the connected bilateral μ-ILED probes. The BLE-based smartphone app was developed using commercial app development software (Android Studio) and used to control the μ-ILED probes for wireless optogenetics.

#### Virus injection and probe implantation surgery

For optogenetic activation of excitatory neurons in the left OFC, we injected the AAV2-CAMKIIα-eYFP (vector biolabs, VB1941) or AAV2-CAMKIIα-ChR2-eYFP (Addgene) virus in the OFC_L_ (AP+2.35, ML±1.3, DV −2.5) of female WT mice between P50 to P60. For optogenetic inactivation of PV^+^ neurons in the left OFC, we injected the AAV9-EF1α-DIO-eYFP (UNC vector core) or AAV9-EF1α-DIO-eArch3.0-eYFP (UNC vector core) virus in the OFC_L_ of female PV-Cre transgenic mice. For the injection, we anesthetized mice using 1.5% isoflurane in oxygen inhalation and performed stereotaxic surgery. Each injection contained 400nl of AAV-infused at a rate of 0.4 nl/sec using a microinjector. We recovered the mice for 7 days after the AAV injection and anesthetized mice again to implant the wireless optogenetic LED probe into the virus-expressing OFC area. Because of the thickness of the probe (0.1mm) and the location of the LED on the probe (+0.22 mm from the tip), we implanted the probe into the anterior and ventral part of the OFC (AP +2.45, ML ±1.3, DV −2.77) to target the LED facing the virus-expressing area without any lesion in it.

#### Behavior test

To connect the small module part to the implanted probe on the mouse head, we briefly anesthetized the mice by isoflurane inhalation. After the module was connected, the mice had 10 min recovery phase in their home cage. On Day0, we habituated mice wearing a module for 15 min in the open field chamber. On Day1, we habituated the mice wearing a module in the sociability chamber for 5 min. We then presented a novel social target and a novel object for 5 min and recorded their approaching behaviors to measure the sociability of the mice. On Day2, we performed a sociability test with blue or green LED stimulation for optogenetic modulation. We gave 20Hz blue light stimulation by 473nm LED for activation or continuous green light stimulation by 532nm LED for inactivation. The LED connected to the probe during the 5 min sociability test session using the custom-designed flexible optoelectronic neural probes. Interaction times of mice were analyzed on Day1 and Day2 as described previously. On Day3, we performed an open field test for 5 min with optogenetic stimulation and 5 min without stimulation to measure the effect of light activation on the locomotion behavior. After the behavior test, we performed the histology of the brain samples from the mice to check the virus expression and probe implanted site. We collected data only if the virus was expressed more than 65% of the total volume of the OFC and the probe was targeted at the virus-expressed area.

### Experimental design & Statistical analysis

All statistical analyses were performed using custom codes in IBM SPSS statistics 21 (IBM) and OriginPro 2019 (Origin). The normal distribution of data was verified using the Shapiro-Wilk normality test. For parametric tests, we used the paired student’s t-test to compare matched pairs, unpaired student’s t-test to compare two groups, the one-way ANOVA with Fisher’s least significant difference [LSD] post hoc test to compare multiple groups, and two-way ANOVA test to compare I-F curves of PV^+^ neurons. For nonparametric tests, we used the Wilcoxon signed-rank test to compare matched pairs, the Mann-Whitney *U* test to compare between two independent groups, and the Kolmogorov–Smirnov test to compare the distributions of two groups. The number of animals analyzed by statistical tests and the exact p values from each experiment were reported within each figure legend.

## Results

### Post-weaning social isolation enhances sociability in female mice

Previous reports have shown that both short- and long-term social isolation affects mouse social behaviors (Makinodan et al., 2012; Matthews et al., 2016). However, the effects differ depending on the duration of isolation, and the data were collected only in male mice. To examine whether short- and long-term social isolation effects on mouse sociability differ between the sexes, we performed the sociability test in both male and female mice. For the sociability test, we put two small chambers at diagonal corners in the maze, each of which contained a novel object (O) or a novel and sex-matched mouse (M), and measured the relative times of mice sniffing or spending at or nearby the novel mouse compared to those times at the novel object as indices of the mouse sociability. We quantified the sociability in mice that had experienced three different post-weaning social conditions at postnatal day 80 (P80): group-housed (GH), socially isolated for one day (1dSI), or socially isolated for 60-days after weaning (post-weaning social isolation, PWSI) (Fig. 1A and 1B). Both the sniffing time of 1dSI mice when they came into contact with a novel mouse and their time spent in the social interaction zone did not differ significantly from those of GH mice in both males and females (Fig. 1C and 1D).

**Fig 1.**
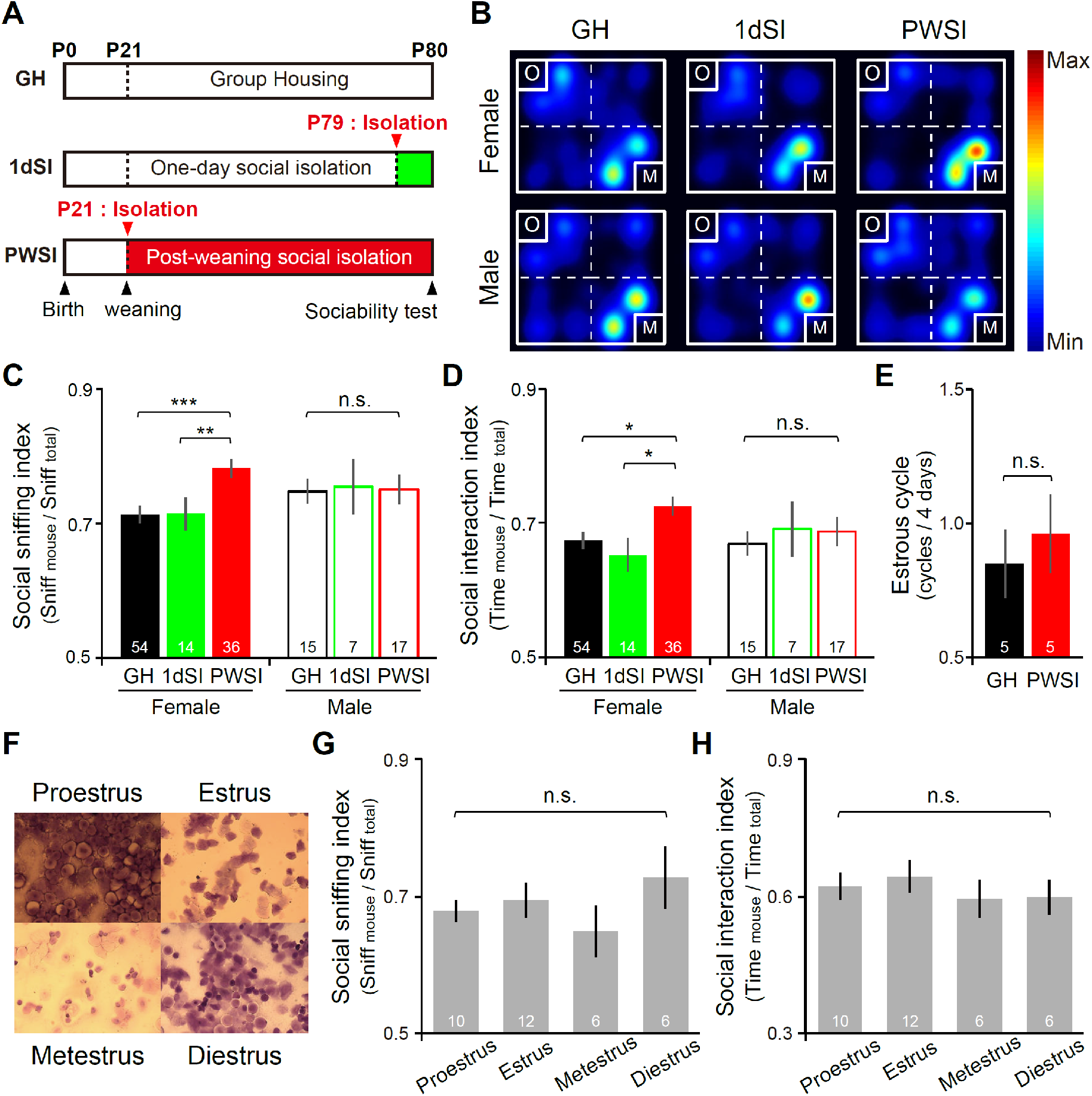
PWSI during adolescence enhances social approaching behavior in female mice. (A) Experimental design of the sociability test. Schematic of group housing (GH), 1day social isolation (1dSI), and post-weaning social isolation (PWSI). Note that the sociability behavior was measured at P80. (B) Representative heatmaps indicating mouse locations in the square chamber during the sociability test. In the chamber, a container containing a novel object (O) and a container containing the same-sex novel mouse (M) were located at diagonal corners. Heatmaps were calculated in EthoVision XT as the mean duration of time spent in an area for each group. (C) The social sniffing index displays the social sniffing times divided by the total sniffing times. One-way ANOVA test, female, F_(2, 101)_ = 8.431, ***p < 0.001; male, F_(2, 36)_ = 0.015, p = 0.985, Fisher’s least significant difference post hoc test (female, GH vs PWSI, ***p <0.001; 1dSI vs PWSI, **p = 0.009; GH vs 1dSI, p = 0.967). Note that only the PWSI female mice showed a significant increase in social sniffing behavior. (D) The social interaction index displays the time spent in the social zone divided by the total time spent in both object and social interaction zones. One-way ANOVA test, female, F_(2, 101)_ = 3.496, *p = 0.034; male, F_(2, 36)_ = 0.109, p = 0.897, Fisher’s least significant difference post hoc test (female, GH vs PWSI, *p = 0.027; 1dSI vs PWSI, *p = 0.031; GH vs 1dSI, p = 0.494). Note that only the PWSI female mice showed a significant increase in social interaction behavior. (E) The number of estrous cycles per 4 days of GH (*n* = 5) and PWSI mice (*n* = 5). The duration of estrous cycles was not different between groups. Mann-Whitney U test, p = 0.912. (F) Representative images of cells in the vaginal smear that were stained with crystal violet. The estrous cycles of female mice were identified based on the shape of stained cells. (G) The social sniffing indices of females across the estrous cycle. One-way ANOVA test, F_(3,30)_ = 0.927, p = 0.440. (H) The social interaction indices of females across the estrous cycle. One-way ANOVA test, F_(3,30)_ = 0.513, p = 0.677. All data are presented as mean ± SEM. For (C-D), Female, GH, *n* = 54 mice, 1dSI, *n* = 14 mice, PWSI, *n* = 36 mice; Male, GH, *n* = 15 mice, 1dSI, *n* = 7 mice, PWSI, *n* = 17 mice; For (G-H), Proestrus, *n* = 10 mice, Estrus, *n* = 12 mice, Metestrus, *n* = 6 mice, Diestrus, *n* = 6 mice. *** p < 0.001, ** p < 0.01, * p < 0.05, n.s., not significant.

Notably, PWSI induced changes in sociability in a sex-specific manner: only female mice showed a significant increase in social sniffing time and social interaction time, while PWSI male mice showed normal sociability as the GH mice (Fig. 1C and 1D). We also confirmed that the effect of PWSI on female mice was not simply due to hormonal changes that occur during the estrous cycle. First, we confirmed that the estrous cycle was not different between GH and PWSI female mice (Fig. 1E). We next confirmed that the mouse sociability (both social sniffing index and social interaction index) that we measured was consistent across the estrous cycle in female mice (Fig. 1F-1H). Therefore, our data indicate that long-term social isolation during adolescence affects females’ social behavior more than males, causing hypersociability in females.

### PWSI reduces PV expression in the OFC_L_ of female mice

We next examined whether the social experience during adolescence is required to mature PV^+^ inhibitory circuits in the OFC. We quantified the number of PV^+^ or PNN^+^ neurons in the OFC of the GH and the PWSI mice at P80. We collected data from male and female mice separately. We first immunostained both for the PV, a marker for fast-spiking GABAergic neurons, and the NeuN, a marker for all neurons (Fig. 2A). We counted the number of immunostained neurons automatically in the three coronal sections that were collected across the center of the OFC (+2.1 ~ +2.34 mm from the bregma) per each hemisphere in each mouse (See Methods; Fig. 2B and 2C). Interestingly, the number of PV^+^ and NeuN^+^ neurons in the OFC was significantly lower in the brains of PWSI female mice compared to the GH female mice, especially in the left hemisphere (Fig. 2D-2F). In male mice, however, the number of PV^+^ and NeuN^+^ neurons was not significantly different between GH and PWSI groups (Fig. 2D-2F). When we counted the number of PV^-^ and NeuN^+^ neurons by subtracting the number of PV^+^ neurons from the total number of NeuN^+^ neurons, there was no difference between groups (Fig. 3F; filled bars; female, OFC_L_, p=1, Mann Whitney U test). These data indicate that the reduction of NeuN^+^ neurons preferentially occurs in the PV^+^ neurons rather than in overall PV^-^ neurons in the OFC_L_ (Fig. 2F). We next quantified the PNN^+^ neurons in the OFC by staining for *Wisteria floribunda agglutinin* (WFA), a marker for PNNs (Fig. 2G). Both PV^+^ and PV^-^ neurons were surrounded by the PNNs in the OFC (Fig. 2H). We found that the numbers of WFA^+^ neurons in the OFC were not different between GH and PWSI in both male and female mice (Fig. 2I). Collectively, our data indicate that social experience during adolescence is crucial for the maturation of PV^+^ neurons, especially the PV expression, in the OFC_L_ of female mice.

**Fig 2.**
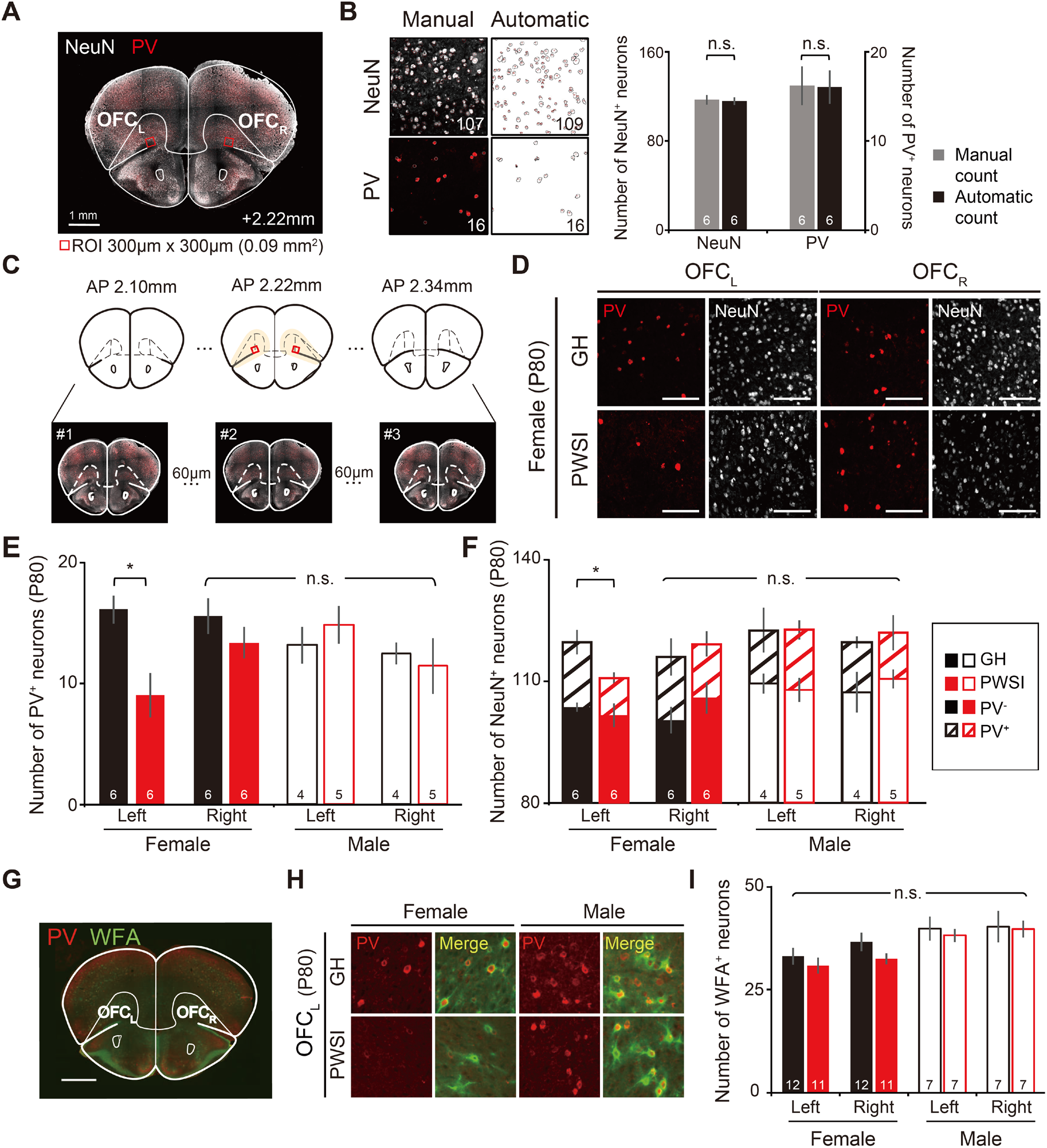
PV expression in the OFC_L_ decreases in the PWSI female mice. (A) A representative image of the immunostained PV and NeuN in the OFC of a mouse brain coronal section marked with the regions of interest (ROI, 0.09 mm^2^). Scale bar, 1 mm. (B) Validation of the automatic cell counting method. Left, representative images of the NeuN (white) and PV (red) immunostained in the OFC_L_ of GH female mice. The left panels show original images used for manual counting, and the right panels show automatically detected cell boundaries from original images. The bottom numbers indicate the counted cell numbers. Right, the average numbers of NeuN^+^ and PV^+^ neurons in the OFC_L_ of GH female mice from the manual counting (gray) and automatic counting (black) (*n* = 6). Note that there is no difference between numbers from manual and automatic counting. Mann-Whitney U test, NeuN, p = 0.200; PV, p = 1.000. (C) Schematic brain images showing the selection of brain slices for cell counting. Three brain slices were selected between AP +2.1mm and +2.34mm with an interval of 60μm. (D) Representative fluorescence images of the PV (red) and the NeuN (white) immunostained in the OFC_L_ and OFC_R_ of P80 GH and PWSI female mice. Scale bars, 100μm. (E) The number of PV^+^ neurons in the OFC. PWSI decreased the number of PV^+^ neurons in the female OFC_L_. Mann-Whitney U test, female left, *p = 0.030; female right, p = 0.297; male left, p = 0.623; male right, p = 1.000. (F) The number of NeuN^+^ neurons in the OFC. Mann-Whitney U test, female left, *p = 0.024, female right, p = 1.000, male left, p = 0.903; male right, p =1.000. Slashed bars indicate the number of PV^+^ (NeuN^+^) neurons and filled bars indicate PV^-^ (NeuN^+^) neurons. Note that, PWSI female mice showed a selective reduction in PV^+^ (NeuN^+^) neurons in OFC_L_. (G) A representative image of a mouse coronal brain section that was immunostained for PV (red) and WFA (green) (scale bar, 1000μm). (H) Representative fluorescence images of the OFC_L_ immunostained with anti-PV and anti-WFA antibodies. (I) The number of WFA^+^ neurons in the OFC left and right of male and female mice. Mann-Whitney U test, GH vs. PWSI, female left, p = 0.46; female right, p = 0.185; male left, p = 0.337; male right, p = 0.442. All data are presented as mean ± SEM. GH, group-housed mice; PWSI, mice isolated from P21 to P80. For (E-F), Female, GH, *n* = 6 mice, PWSI, *n* = 6 mice; Male, GH, *n* = 4 mice, PWSI, *n* = 5 mice. For (H-J), Female, GH, *n* = 12 mice, PWSI, *n* = 11 mice; Male, GH, *n* = 7 mice, PWSI, *n* = 7 mice. *p < 0.05, n.s., not significant.

**Fig 3.**
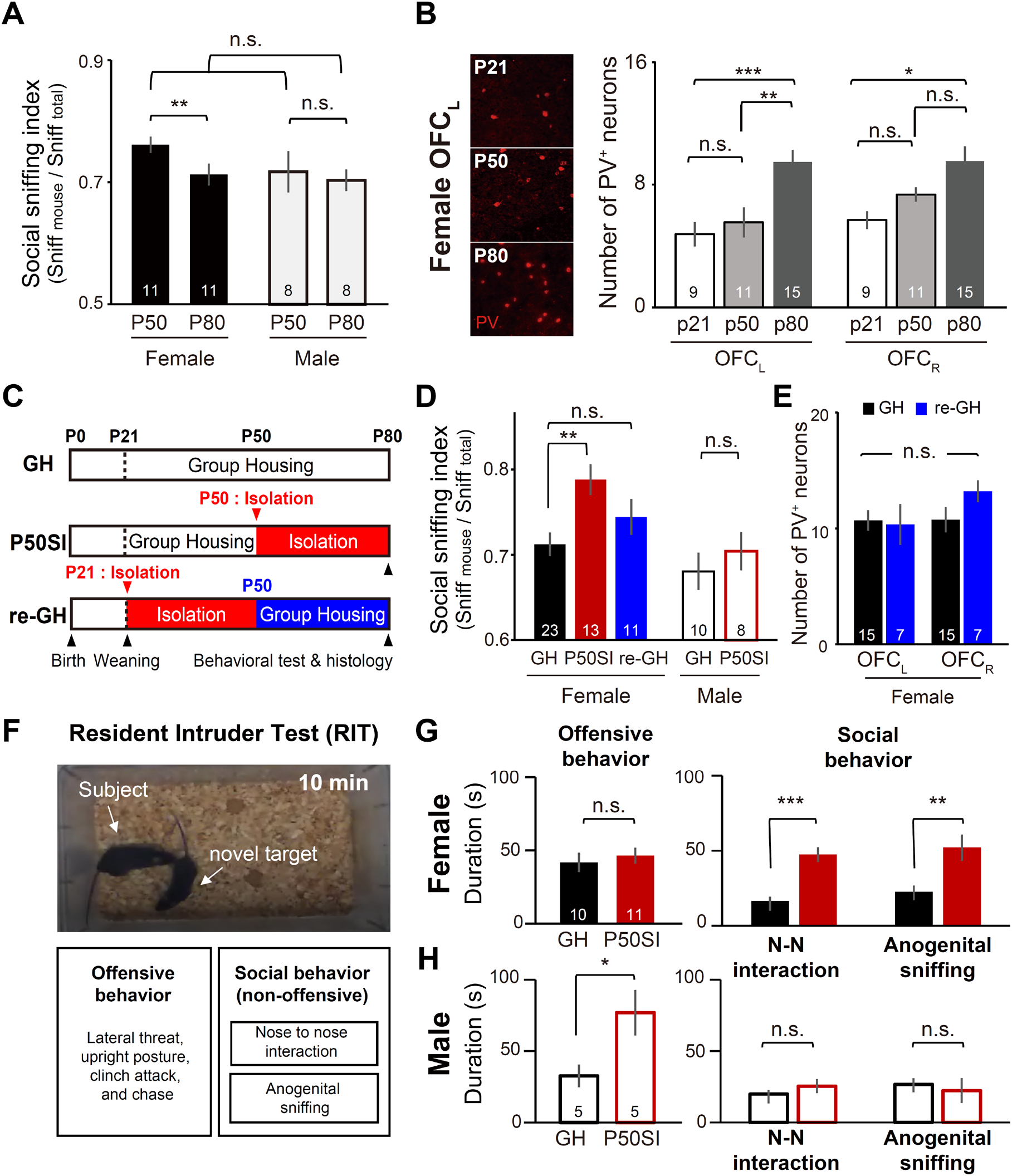
Social isolation during late adolescence causes hypersociability only in female mice. (A) The social sniffing indices of GH female and male mice. The sociability test was performed at P50 and P80 in the same mouse. Paired, Student’s t-test within a group, female, **p = 0.008; male, p = 0.756. Unpaired, Student’s t-test between groups, P50, p = 0.285; P80, p = 0.731. Female, *n* =11 mice; Male, *n* =8 mice. (B) Left, representative fluorescence images of the OFC_L_ immunostained with anti-PV antibodies (red). Brain sections were collected from P21 (top), P50 (middle), and P80 (bottom) GH female mice. Right, the number of PV^+^ neurons in the OFC_L_ and OFC_R_ of female mice. One-way ANOVA test, female OFC_L_, F_(2, 32)_ = 8.473, **p=0.001, Fisher’s least significant difference post hoc test (P21 vs P50, p = 0.580; P50 vs P80, **p = 0.003; P21 vs P80, ***p < 0.001); female OFC_R_, F_(2, 28)_ = 2.856, *p = 0.010, Fisher’s least significant difference post hoc test (P21 vs P50, p = 0.204; P50 vs P80, p = 0.065; P21 vs P80, **p = 0.003). Female, P21, *n* = 9 mice; P50, *n* = 11 mice; P80, *n* = 15 mice. (C) Schematic of group-housing (GH), social isolation from P50 to P80 (P50SI), and re-group-housing at P50 after social isolation at P21 (re-GH). Note that the behavioral tests were performed at P80. (D) The social sniffing indices of GH, P50SI, and re-GH mice. One-way ANOVA test, female, F_(2, 44)_ = 5.069, *p = 0.01, Fisher’s least significant difference post hoc test (female, GH vs P50SI, **p = 0.007; GH vs re-GH, p = 0.276; P50SI vs re-GH, p = 0.127). Mann-Whitney U test, male, GH vs P50SI, p = 0.505. Note that only the P50SI female mice showed a significant increase in social sniffing behavior compared to GH female mice. Female, GH, *n* = 23 mice, P50SI, *n* = 13 mice, re-GH, *n* = 11 mice; Male, GH, *n* = 10 mice, P50SI, *n* = 8 mice. (E) The number of PV^+^ neurons in the OFC of GH (black) and re-GH (blue) female mice. Unpaired, Student’s t-test, OFC left, p = 0.873; OFC right, p = 0.120. Female, GH, *n* = 15 mice; re-GH, *n* = 7 mice. (F) The resident-intruder test (RIT) was performed for P50SI and GH female and male mice at P80. Top, a representative image of RIT. Bottom, the list of analyzed behaviors. The offensive behavior included lateral threat, upright posture, clinch attack, and chase. The social behavior included nose to nose interaction (N-N interaction) and anogenital sniffing. (G) The RIT results of GH and P50SI female mice. Left, duration of offensive behaviors. Mann-Whitney U test, p = 0.699. Right, duration of social behaviors. Mann-Whitney U test, N-N interaction, ***p < 0.001; anogenital sniffing, **p = 0.008. Female P50SI mice showed a significant increase in social behaviors without any change in offensive behaviors. (H) The RIT results of GH and P50SI male mice. Left, duration of offensive behaviors. Mann-Whitney U test, *p = 0.037. Right, duration of social behaviors. Mann-Whitney U test, N-N interaction, p = 0.095; anogenital sniffing, p = 0.835. Male P50SI mice showed a significant increase in offensive behaviors without any change in social behaviors. All data are presented as mean ± SEM. For (F-H), Female, GH, *n* = 11 mice, P50SI, *n* = 11 mice; Male, GH, *n* = 5 mice, P50SI, *n* = 5 mice. ***p < 0.001, **p < 0.01, *p < 0.05, n.s., not significant.

### Late adolescence is a critical window for maturing sociability in female mice

PWSI mice experienced long-term social isolation from when they were weaned at P21 to when they became adults at P80. This period includes both early juvenile periods before the onset of puberty and late adolescence after the onset of puberty. Since puberty leads to sexually dimorphic changes in behaviors, we wondered how mouse sociability matures after the onset of puberty in males and females. We thus tracked sociability changes from P50 (late adolescence (Laviola et al., 2003) to P80 (adult) in both sexes of GH mice by measuring the social sniffing index at P50 and at P80. Interestingly, we found a significant decrease in sociability at P80 in female mice but not in male mice (Fig. 3A). We further found that during this period, PV^+^ neurons increased significantly in female OFC_L_. In contrast, OFC_R_ did not show such an increase between P50 and P80 period (Fig. 3B). These data suggest that late adolescence from P50 to P80 is a critical window for the maturation of sociability with an increase in PV^+^ neurons in the OFC_L_ in female mice.

We next examined the effect of social isolation during late adolescence in both female and male mice. We directly isolated the mice after P50 until P80 (P50SI; Fig. 3C). In addition, we re-group-housed a group of PWSI female mice at P50 to make them re-experience social interaction during late adolescence (re-GH; Fig. 3C). The P50SI female mice showed an increase in sociability like the PWSI female mice, while reGH female mice nor P50SI male mice did not show any changes in sociability compared to control GH mice (Fig. 3D). Not only our modified 3-chamber-like test, but a standard 3-chamber test also showed an increase in sociability in P50SI female mice (data not shown). In addition, the number of PV^+^ neurons in re-GH female mice was not significantly different from that in GH female mice in both hemispheres of the OFC (Fig. 3E), indicating that social experience during late adolescence was sufficient to mature the PV^+^ neurons in the OFC_L_.

Finally, we wondered whether the enhanced sociability in P50SI female mice, measured by our sociability test, was related with changes in aggression. To examine this, we performed the resident-intruder test (Tan et al., 2021) on P50SI male and female mice. We counted both offensive behaviors, such as threatening and attacking behaviors, and social behaviors, such as nose sniffing and anogenital sniffing, separately during the test (Fig.3F). In female mice, the aggression level did not increase by P50SI. Still, the social approaching behaviors significantly increased (Fig. 3G). On the other hand, P50SI male mice showed a significant increase in aggression without any changes in social behaviors (Fig. 3H). Collectively, our data indicate that social isolation during late adolescence induced hypersociability in female mice without increasing aggression whereas it induced hyperaggression in male mice.

### PWSI disrupts social sniffing-induced activity in the OFC_L_ of female mice

Because PV^+^ neurons in the OFC_L_ were found to be less numbered following PWSI in female mice, we investigated whether the OFC_L_ neural response to social contact has been altered in head-fixed female mice by *in vivo* multichannel recordings (Fig. 4A and 4B). During the recording, both GH and PWSI female mice showed sniffing behaviors toward an object or another mouse under the head-fixed condition, and the number and duration of sniffing times did not differ between groups (Fig. 4C and 4D). We identified neurons that showed significant sniffing-induced activity in the OFC (Fig. 4E-4H). Interestingly, in GH females, more neurons showed sniffing-induced activity in the OFC_L_ than in the OFC_R_ responses, and their responses were stronger when a mouse sniffed another mouse than when it sniffed an object (Fig. 4E and 4F). More importantly, these socially responsive neurons in the OFC_L_ of female mice largely disappeared after PWSI (Fig. 4E, 4G, and 4H). On the other hand, the sniffing-induced activity of the OFC_R_ did not differ between GH and PWSI female mice (Fig. 4F). Among the responsive neurons in the OFC_L_, the fraction of the fast-spiking neurons was smaller in the PWSI mice than in the GH mice. These data indicate that PWSI decreased social sniffing-induced activity in the OFC_L_ neurons, particularly including fast-spiking neurons, and reduced overall social representation in the OFC_L_.

**Fig 4.**
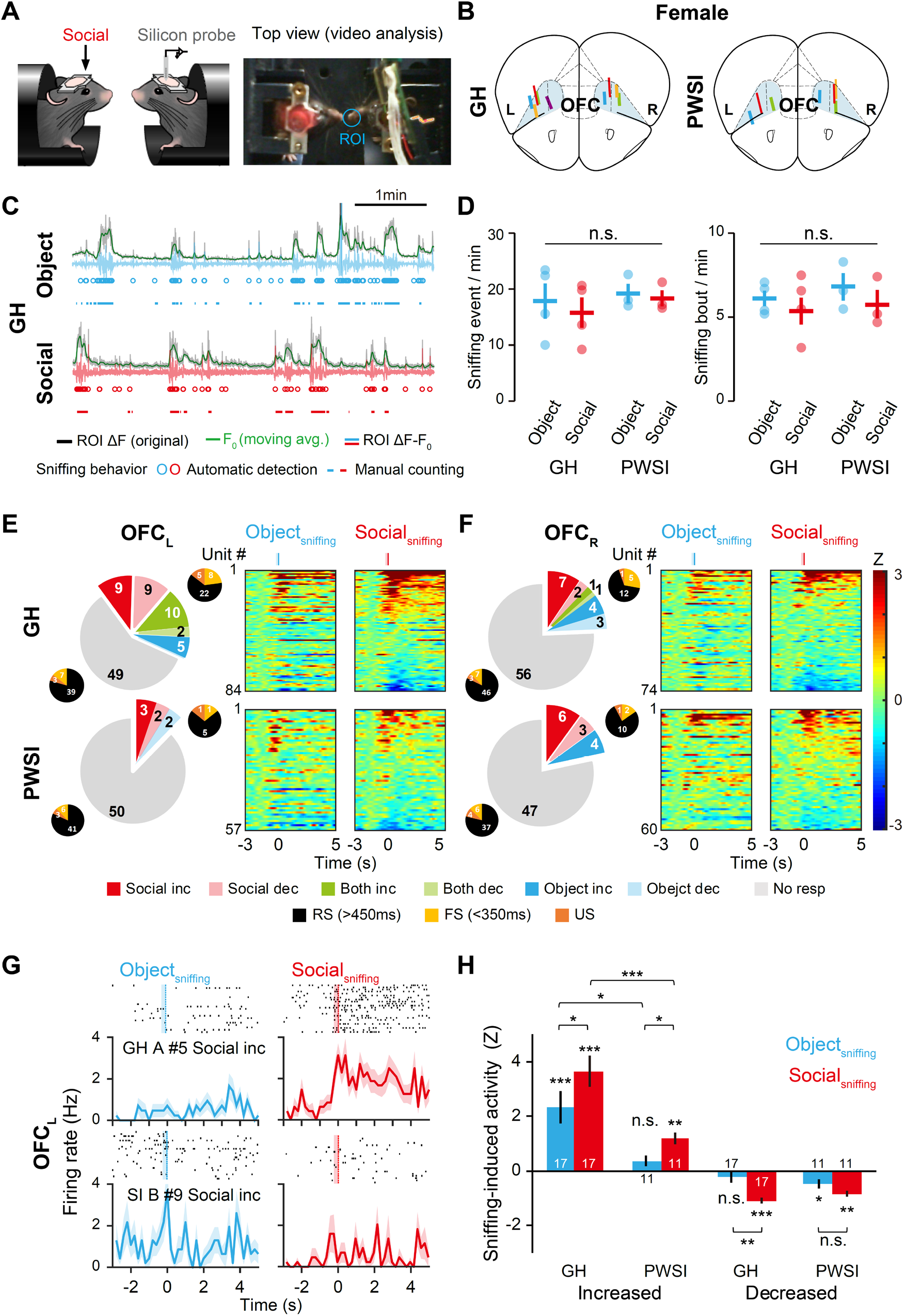
PWSI reduces sniffing-induced activity in the OFC_L_ of female mice. (A) Left, experimental design for *in vivo* multichannel recording of the neurons responsive to social sniffing in the OFC of head-fixed mice. Right, a representative top view image for video analysis. The blue circle indicates the region of interest (ROI) used to analyze the brightness changes in the video clip. (B) Schematic illustration of mouse brain sections marked with the recording sites in the OFC of GH or PWSI female mice. Blue shading, OFC; colored bars, recording sites confirmed by post hoc histology. (C) Representative sniffing events measured in GH female mice. Gray trace, raw brightness changes in the ROI (ROI ΔF). Green trace, moving average in the ROI (F_0_, sliding window = 50 frames). Blue and red trace, normalized brightness changes in the ROI (ROI ΔF–F_0_). Circles, sniffing events were automatically detected by a Matlab-based image analysis code. Horizontal lines, sniffing events manually detected by an observer in the EthoVision XT software. Note that automatic counting methods detect sniffing events at similar times identified by manual counting. Blue, object sniffing; red, social sniffing. (D) Left, sniffing events automatically counted. Dots, individual data; lines, means ± SEM. Wilcoxon signed-rank test within a group, Object vs. Social, GH, p = 0.465, PWSI, p = 1.000. Mann-Whitney U test between groups, GH vs. PWSI, Object, p = 0.857, Social, p = 0.629. Right, the sniffing events manually identified. Dots, individual data; lines, means ± SEM. Wilcoxon signed-rank test within a group, Object vs. Social, GH, p = 0.686, PWSI, p = 0.109. Mann-Whitney *U* test between groups, GH vs. PWSI, Object, p = 0.393, Social, p = 1.000. GH, *n* = 4 mice, PWSI, *n* = 3 mice. (E) Neural responses associated with sniffing in the female OFC_L_. The large pie charts, the OFC_L_ neurons showing social sniffing responses (Social, red), object sniffing responses (Object, blue), or both social and object sniffing responses (Both, green). Inc and dec, neurons showing significantly increased and decreased activities after the onset of sniffing, respectively (p < 0.01, bootstrap); No resp., neurons that show no responses. The small pie charts, the waveforms of the responsive (right-top) and non-responsive (left-bottom) OFC_L_ neurons classified according to the waveform width. RS, regular-spiking neurons with a waveform width >450ms; FS, fast-spiking neurons with a waveform width <350ms; US, unclassified neurons with a waveform width between 350ms and 450ms. Heatmaps, *Z*-scored activity of all neurons normalized by the pre-sniffing activity. GH, *n* = 5 mice, *n* = 84 units; PWSI, *n* = 3 mice, *n* = 57 units. (F) The same as (C) but for the OFC_R_ neurons. GH, *n* = 4 mice, *n* = 74 units; PWSI, *n* = 4 mice, *n* = 60 units. (G) Representative raster plots and peri-event time histograms of example ‘Social Inc’ OFC_L_ neurons from the GH female mouse (top, GH A #5) and from the PWSI female mouse (bottom, SI B #9) aligned by the onset of object sniffing (left, blue) and the onset of social sniffing (right, red). (H) Averaged sniffing-induced activity for 2 sec (*Z* score) of the top 20% of OFC_L_ neurons that show either increased or decreased responses after the onset of social sniffing. Bars, means ± SEM. Wilcoxon signed-rank test for each bar, Object_sniffing_, Social_sniffing_, ***p < 0.001, ***p < 0.001 for GH Increased; p = 0.083, **p = 0.004 for PWSI Increased; p = 0.478, ***p < 0.001 for GH Decreased; *p = 0.045, **p = 0.004 for PWSI Decreased. Wilcoxon signed-rank test within a group, Object_sniffing_ vs. Social_sniffing_, *p = 0.047 for GH Increased; *p = 0.023, for PWSI Increased; **p = 0.002 for GH Decreased; p = 0.120 for PWSI Decreased; Mann-Whitney U test between groups, Object_sniffing_, Social_sniffing_, *p = 0.019, ***p < 0.001 for Increased activity; p = 0.452, p = 0.110 for Decreased activity. GH, *n* = 17 units, PWSI, *n* = 11 units. ***p < 0.001, **p < 0.01, *p < 0.05, n.s., not significant.

### PV^+^ expression, not PNNs in the OFC_L_, shapes social behavior in female mice

As we found that PWSI impaired the PV expression in the OFC_L_ and increased social behavior, we next examined the causal relationship between the reduction of PV expression in the OFC_L_ and the increase in the sociability of female mice. To reduce the PV expression *in vivo*, we used the CRISPR/Cas9 system (Swiech et al., 2015) and designed two pairs of single-guide RNAs (sgRNAs) to target the 4^th^ and 5^th^ exons of the PV gene (sgPV). We first confirmed that those sgRNAs knocked out the PV gene by either deletion or insertion of the targeted sequences in more than 85% of the Cas9-transfected Neuro2a cells (Fig. 5A). We then constructed the adeno-associated viral vectors (AAVs) to deliver those sgRNAs together with the tdTomato gene to neurons *in vivo* (Fig. 5A) and injected the AAVs into the cortex of PV::Cas9-eGFP mice, which were created by crossing PV-Cre with Cas9^fl/fl^-eGFP mice (sgPV; and Fig. 5B-C). As a control, we injected the AAV that expressed only the tdTomato without sgRNAs into the crossed mice (tdTom). In the sgPV mice, originally PV^+^ neurons were expressed with both Cas9-eGFP and tdTomato, and their PV gene was knocked out by Cas9 in the PV^+^ neurons selectively (Fig. 5B-C).

**Fig 5.**
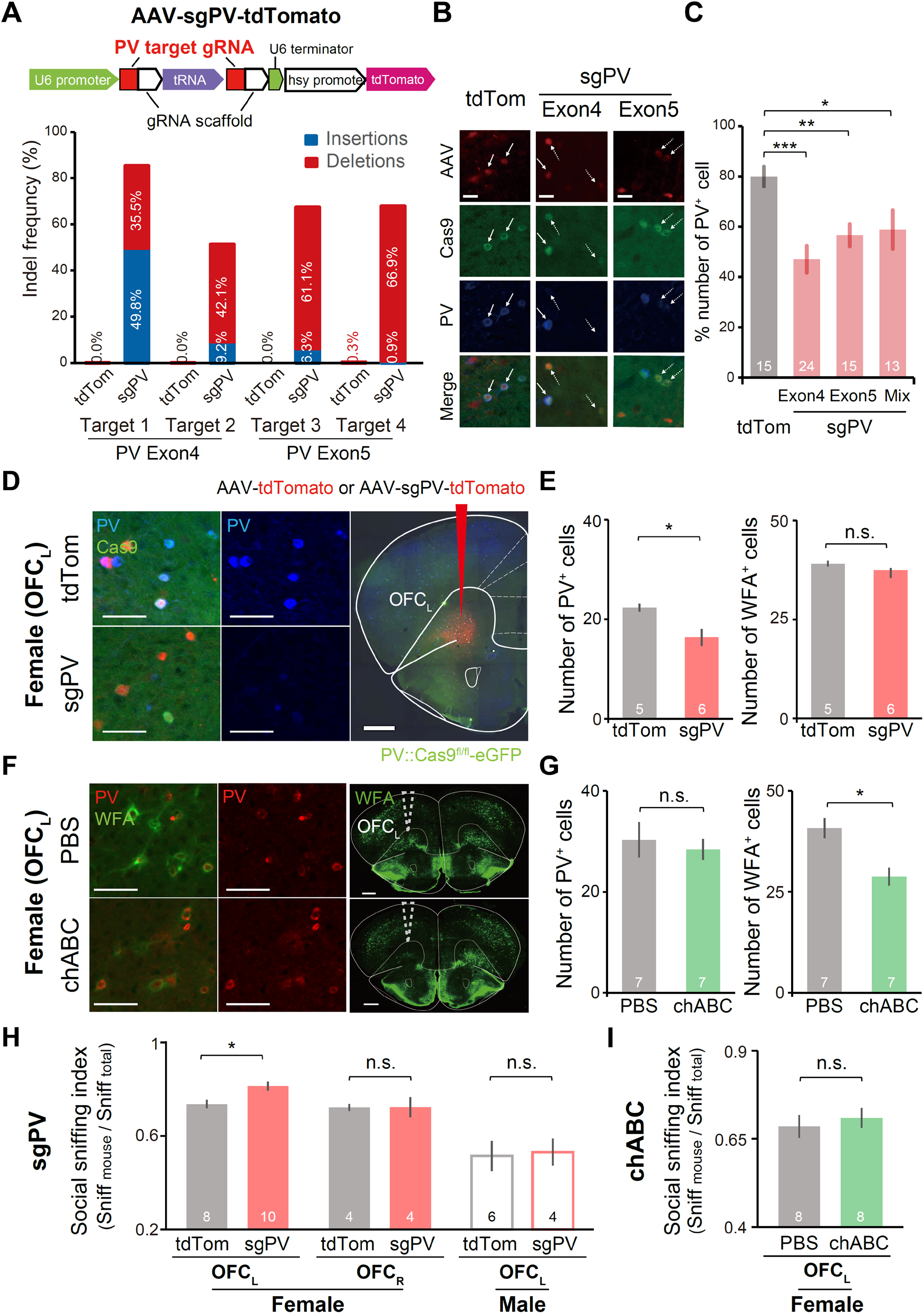
CRISPR/Cas9-mediated knock-down of PV, but not PNN degradation, in the OFC_L_ enhances sociability in female mice. (A) Top, schematic illustration of the AAV-sgPV-tdTomato expression vector. The sgRNA vector contains the encoding sequence of tdTomato protein for identifying transduced neurons. We used two types of AAVs to target both exon 4 and exon5 of the PV gene. Bottom, DNA insertion (blue) and deletion (red) mutation ratio (Indel frequency) by the designed sgPVs targeting exon 4 and exon5 of the PV gene and cloned in the AAV-tdTom vector. (B) Representative PV-immunostained images of PV::Cas9 mice injected with AAV-tdTomato or AAV-sgPV-tdTomato. Scale bars, 20μm. Red, tdTomato expressed by AAV vectors; green, Cas9-eGFP transgenically expressed in PV^+^ neurons; blue, immunostained PV proteins. Solid arrows indicate triple-positive cells, and dashed arrows indicate double-positive and PV-negative cells. (C) Quantification of PV^+^ cells among eGFP and tdTomato double-positive cells in the retrosplenial cortex (RSC), injected with AAVs that deliver tdTom or sgPVs, compared to the Cas9-eGFP expression in PV::Cas9 mice. Numbers in columns indicate the number of brain samples analyzed. One-way ANOVA, F_(3, 63)_ = 6.478, ***p < 0.001; Fisher’s least significant difference post hoc test. (D) CRISPR/Cas9-mediated knock-down of PV in the female OFC_L_. Left and middle, representative fluorescence images of the OFC_L_ of the female PV::Cas9-eGFP mice injected with the AAV-tdTomato (top, tdTom) or AAV-sgPV-tdTomato (bottom, sgPV). Red, tdTomato; green, Cas9-eGFP; blue, immuno-stained PV. Scale bars, 50μm. Right, a representative image of the OFC_L_ of a PV::Cas9-eGFP mouse injected with the AAV-sgPV-tdTomato. Scale bar, 1000μm. (E) The number of PV^+^ and WFA^+^ neurons in the OFC_L_ injected with the AAV-tdTomato (gray bars) or with the AAV-sgPV-tdTomato (red bars). Mann-Whitney U test, PV, *p = 0.022; WFA, p = 0.306. Female, *n* = 5 mice for tdTom, *n* = 6 mice for sgPV. (F) chABC-induced PNN degradation in the OFC_L_ of female mice. Left and middle, immunostained PV (red) and WFA (green) in the OFC_L_ of female mice injected with PBS (top) or chABC (bottom). Scale bars, 50μm. Right, representative fluorescence images of the PNN stained with the WFA Note that the chABC injection selectively degraded PNNs in the OFC_L_. Scale bars, 1000μm. (G) The number of PV^+^ and WFA^+^ neurons in the OFC_L_ injected with the PBS (gray bars) or the chABC (green bars). Mann-Whitney U test, PV, p = 0.609; WFA, *p = 0.017. Female, *n* = 7 mice for PBS, *n* = 7 mice for chABC (H) Social sniffing indices of PV::Cas9 female mice injected with AAV-tdTomato (tdTom) or AAV-sgPV-tdTomato (sgPV). Mann-Whitney U test, tdTom vs sgPV, female OFC_L_, *p = 0.021; female OFC_R_, p = 0.885; male OFC_L_, p = 0.914. tdTom, female OFC_L_, *n* = 8 mice; female OFC_R_, *n* = 4 mice, male OFC_L_, *n* = 6 mice; sgPV, red, female OFC_L_, *n* = 10 mice; female OFC_R_, *n* = 4 mice, male OFC_L_, *n* = 4 mice. (I) Social sniffing indices of mice injected with PBS (gray) or chABC (green). Mann-Whitney U test, p = 0.878. PBS, *n* = 8 mice; chABC, *n* = 8. All data are presented as mean ± SEM. *p < 0.05, n.s., not significant.

To knock down PV in the OFC_L_ during late adolescence until mice became adults, we injected the AAVs into the OFC_L_ when mice were at around P50 and examined changes in the sociability of the injected mice at P80. We confirmed the reduction of PV expression in the eGFP^+^/tdTomato^+^ neurons without any decrease in PNN expression by immunostaining the PV and the PNNs in the OFC_L_ (Fig. 5D-E). The female mice with PV knock-down in the OFC_L_ showed a significant increase in sociability (Fig. 5H), similar to the PWSI or P50SI females (Fig. 1B-1D and Fig. 3D). When we knocked down the PV in the OFC_R_ of female mice or the OFC_L_ of male mice, we did not see such an increase in sociability compared to the control mice (Fig. 5H). We further examined whether a reduction in PNNs surrounding PV^+^ neurons affects female sociability. We infused chondroitinase ABC (chABC), an enzyme that degrades PNNs by removing sulfate side chains of proteoglycans, into the OFC_L_ (Fig. 5F). Treatment with chABC significantly decreased the number of WFA^+^ neurons in the OFC_L_ without affecting PV expression (Fig. 5G). Not like the female mice knocked down with PV, the female mice treated with chABC showed normal sociability (Fig. 5I). Together, our data indicate that the PV expression, but not the PNN maturation, in the OFC_L_ was a key factor shaping the social behavior of female mice.

### Decreased PV expression reduces social sniffing-induced activity via hyper-excitation of OFC_L_ neurons

We next investigated whether the decrease of PV expression in the OFC_L_ was responsible for the reduced sniffing-induced activity in the OFC_L_ of the PWSI female mice. We performed *in vivo* recordings in female mice injected with sgPV AAVs (**sgPV**) or injected with tdTomato AAVs (**tdTom**) in their OFC_L_ (Fig. 6A-6G). First, we found that the average baseline firing rates were significantly higher in the spPV group compared to the tdTom group (Fig. 6D and 6E). Furthermore, similar to PWSI, knocking down PV by sgPV decreased social sniffing-induced activity in the OFC_L_ of female mice (**sgPV**; Fig. 6F and 6G). Conversely, the expression of tdTomato in the OFC_L_ did not alter the sniffing-induced activity of female mice (**tdTom**; Fig. 6F and 6G). Knocking down PV in the OFC_L_ did not affect the sniffing behavior itself, as the number and duration of sniffs in the altered mice were similar to those in the tdTom control mice (Fig. 6C). These data indicate that the knock-down of PV increased overall firing activity while reducing the sniffing-induced activity in the OFC_L_.

**Fig 6.**
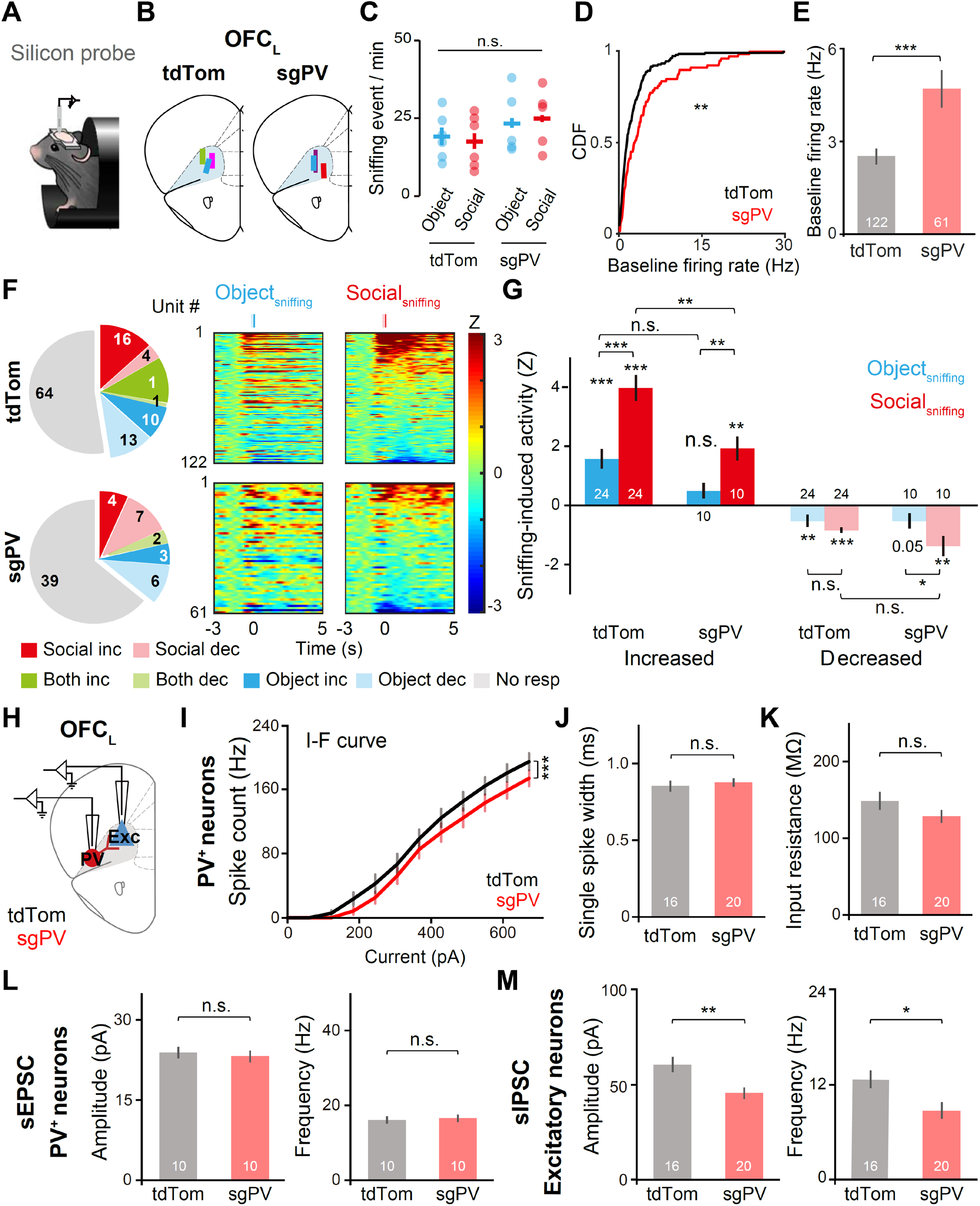
Knock-down of PV reduces sniffing-induced activity and decreases synaptic inhibition in the female OFC_L_. (A) Schematic illustration of *in vivo* multichannel recording in a head-fixed mouse. (B) Schematic illustration of coronal brain sections of *in vivo* recording sites in the OFC_L_ where tdTomato (tdTom)- or PV sgRNA (sgPV)-expressing AAVs were injected. (C) Sniffing events automatically counted for tdTom and sgPV mice. Left, Wilcoxon signed-rank test for within a group, tdTom, p = 0.463; sgPV, p = 0.892. Mann-Whitney U test between groups, Object, p = 0.537; Social, p = 0.126. (D) The cumulative distribution function of baseline firing rates in tdTom and sgPV mice. Baseline firing rates were calculated as the mean firing rates for 2 sec before the sniff onset. Kolmogorov– Smirnov test, **p = 0.006. (E) Mean baseline firing rates of tdTom and sgPV mice. Mann-Whitney U test, ***p < 0.001. (F) Neural responses associated with the sniffing in the OFC_L_ of tdTom- (top) or sgPV-(bottom) injected PV::Cas9 female mice. Pie charts, the OFC_L_ neurons showing social sniffing responses (Social, red), object sniffing responses (Object, blue), or both social and object sniffing responses (Both, green). Inc and dec, neurons showing significantly increased and decreased activities after the onset of sniffing, respectively (p < 0.01, bootstrap); No resp, neurons that show no responses. Heatmaps, Z-scored activity of all recorded neurons by the pre-sniffing activity. Note that the number of sniffing-responsive neurons and social sniffing-induced activity was reduced in sgPV females. Heat maps, *Z*-scored activity of all neurons normalized by the pre-sniffing activity. (G) Averaged sniffing-induced activity for 2 sec (*Z* score) of the top 20% of OFC_L_ neurons that show either increased or decreased responses after the onset of social sniffing. Wilcoxon signed-rank test for each bar, Object_sniffing_, Social_sniffing_, ***p < 0.001, ***p < 0.001 for tdTom Increased; p = 0.154, **p = 0.006 for sgPV Increased; **p = 0.007, ***p < 0.001 for tdTom Decreased; p = 0.053, **p = 0.006 for sgPV Decreased. Note that the increased neurons showed larger activity changes after social sniffing than after object sniffing. Wilcoxon signed-rank test within a group, Object_sniffing_ vs Social_sniffing_, ***p < 0.001 for tdTom Increased; **p = 0.008 for sgPV Increased; p = 0.141 for tdTom Decreased; *p = 0.011 for sgPV Decreased. The social sniffing-induced activity was larger in tdTom females than in sgPV females. Mann-Whitney U test between groups, Object_sniffing_, Social_sniffing_, p = 0.067, **p = 0.004 for Increased activity; p = 0.720, p = 0.073 for Decreased activity. tdTom, *n* = 24 units, sgPV, *n* = 12 units. (H) Physiological properties of PV^+^ neurons expressed with tdTom or sgPV. Schematic illustration of the whole-cell patch-clamp recordings of PV^+^ neurons (red) or excitatory neurons (blue) in the PV::Cas9 mice injected with the AAVs expressing either tdTom or sgPV-tdTom into the OFC_L_. (I) SgPV expression decreased the excitability of PV^+^ neurons. Current-firing (I-F) curves measured in the OFC_L_ PV^+^ neurons. sgPV expression caused a significant decrease in current-evoked spike counts in the PV^+^ neurons. Two-way ANOVA, F_AAV (1, 372)_ = 14.528, ***p < 0.001, Fcurrent (11, 372) = 121.158, ***p < 0.001, F_AAV x current (11, 372)_ = 0.388, p = 0.960. (J) Mean half-width of spikes (ms) measured in PV^+^ neurons expressed with the tdTom or the sgPV. Unpaired, Student’s t-test, p = 0.628. (K) Mean input resistance (MΩ) measured in PV^+^ neurons expressed with the tdTom or the sgPV. Unpaired, Student’s t-test, p = 0.173. (L) Spontaneous EPSCs measured in PV^+^ neurons of the OFC_L_. Left bars, mean amplitudes (pA) of sEPSCs in PV^+^ neurons. Unpaired, Student’s t-test, p = 0.636. Right bars, mean frequency (Hz) of sEPSCs in PV^+^ neurons. Unpaired, Student’s t-test, p = 0.940. tdTom = 10 cells (5 mice); sgPV=10 cells (4 mice); female. (M) Spontaneous IPSCs measured in excitatory neurons of the OFC_L_. Left bars, mean amplitudes (pA) of sIPSCs in Pyr neurons. Unpaired, Student’s t test, **p = 0.006. Right bars, mean frequencies (Hz) of sIPSCs in Pyr neurons. Unpaired, Student’s t test, *p = 0.015. tdTom = 16 cells (3 mice); sgPV=17 cells (3 mice); female. All data are presented as mean ± SEM. For (D-G), tdTom: *n* = 6 mice, *n* = 122 units, sgPV: *n* = 5 mice, *n* = 61 units. For (I-K), tdTom = 16 cells (3 mice); sgPV = 20 cells (4 mice); female. ***p < 0.001, **p < 0.01, *p < 0.05, n.s., not significant.

As we observed an increase in the baseline activity of OFC_L_ neurons by knocking down the PV, we next examined whether the knock-down of PV has altered the excitation-inhibition balance in the OFC_L_ of female mice. We first performed patch-clamp recordings of the PV^+^ neurons in the OFC_L_ by targeting tdTomato^+^ neurons of PV::Cas9 mice injected with sgPV or tdTom AAVs into the OFC_L_ (Fig. 6H-6L). We found that the knock-out of the PV gene by sgPV in PV^+^ neurons reduced the intrinsic excitability of those neurons (Fig. 6I) without any changes in spontaneous excitatory synaptic currents (sEPSCs; Fig. 6L). Other membrane properties, such as the half-width of spike waveforms and the input resistance, were normal in those neurons (Fig. 6J and 6K). We next measured the spontaneous inhibitory postsynaptic currents (sIPSCs) in the OFC_L_ neurons, which were not labeled with the tdTomato in the sgPV and the tdTom mice. These neurons were regular spiking and thus putative excitatory neurons in the OFC_L_ (Fig. 6H and 6M). The OFC_L_ excitatory neurons in the sgPV mice showed a significant decrease in both amplitudes and frequencies of spontaneous IPSCs compared to the tdTom control mice (Fig. 6M). These data indicate that the decreased expression of PV in the OFC_L_ reduced the excitability of the PV^+^ neurons and, in turn, reduced inhibition of the neighboring neurons in the OFC_L_ of female mice.

Our recording data suggested that the decrease in inhibition upon knocking down the PV gene caused an increase in excitation, which led to a rise in the baseline activity of OFC_L_ and disruption in sniffing-induced activities (Fig. 6D-6G). We thus examined whether the abnormal increase of OFC_L_ activity caused hypersociability in female mice. We expressed the channelrhodopsin-2 (ChR2) in the excitatory neurons of the OFC_L_ by injecting the AAV virus, which expresses the ChR2 fused to the eYFP under the CaMKIIα promotor, to the wild-type female mice (Fig. 7A and 7C). We then delivered the light stimulation to the OFC_L_ by inserting a wireless optogenetic device that we developed (Kim et al., 2021; Qazi et al., 2021) to provide the 20Hz blue lights during the sociability tests (Fig. 7B; see Methods). The stimulation was given on the second day of the sociability tests. We measured the first-day sociability (Day1) without the light stimulation and did not observe any differences in the sociability between the ChR2-eYFP and the eYFP groups. We tested their sociability again on the second day with the blue-light stimulation on the OFC_L_ (Day2). The eYFP control mice showed a significant decrease in their sociability on Day2 compared to Day1, potentially due to habituation (Fig. 7D and E). On the other hand, the ChR2-eYFP-expressed mice showed higher levels of social sniffing behaviors with light stimulation than the control mice on Day2 (Fig. 7D). This data indicates that the optogenetic activation of the OFC_L_ induced higher sociability and almost no habituation in social approaching behaviors on Day2 (Fig. 7D and 7E). We next optogenetically inactivated the PV^+^ neurons in OFC_L_ directly during the sociability test on Day2 by expressing the Archaerhodopsin-3 (eArch3.0) in PV^+^ neurons and delivering 532nm light through our wireless probe (Fig. 7G). Similar to the optogenetic activation of excitatory neurons in the OFC_L_, inhibition of PV^+^ neurons induced hypersociability in female mice without habituation on Day 2, when we gave the light stimulation in the OFC_L_ (Fig. 7H and 7I). The light stimulation did not affect the locomotion in both groups (Fig. 7F and 7J). Collectively, our data demonstrate that hyper-excitation of the OFC_L_ by reducing inhibition caused hypersociability in female mice.

**Fig 7.**
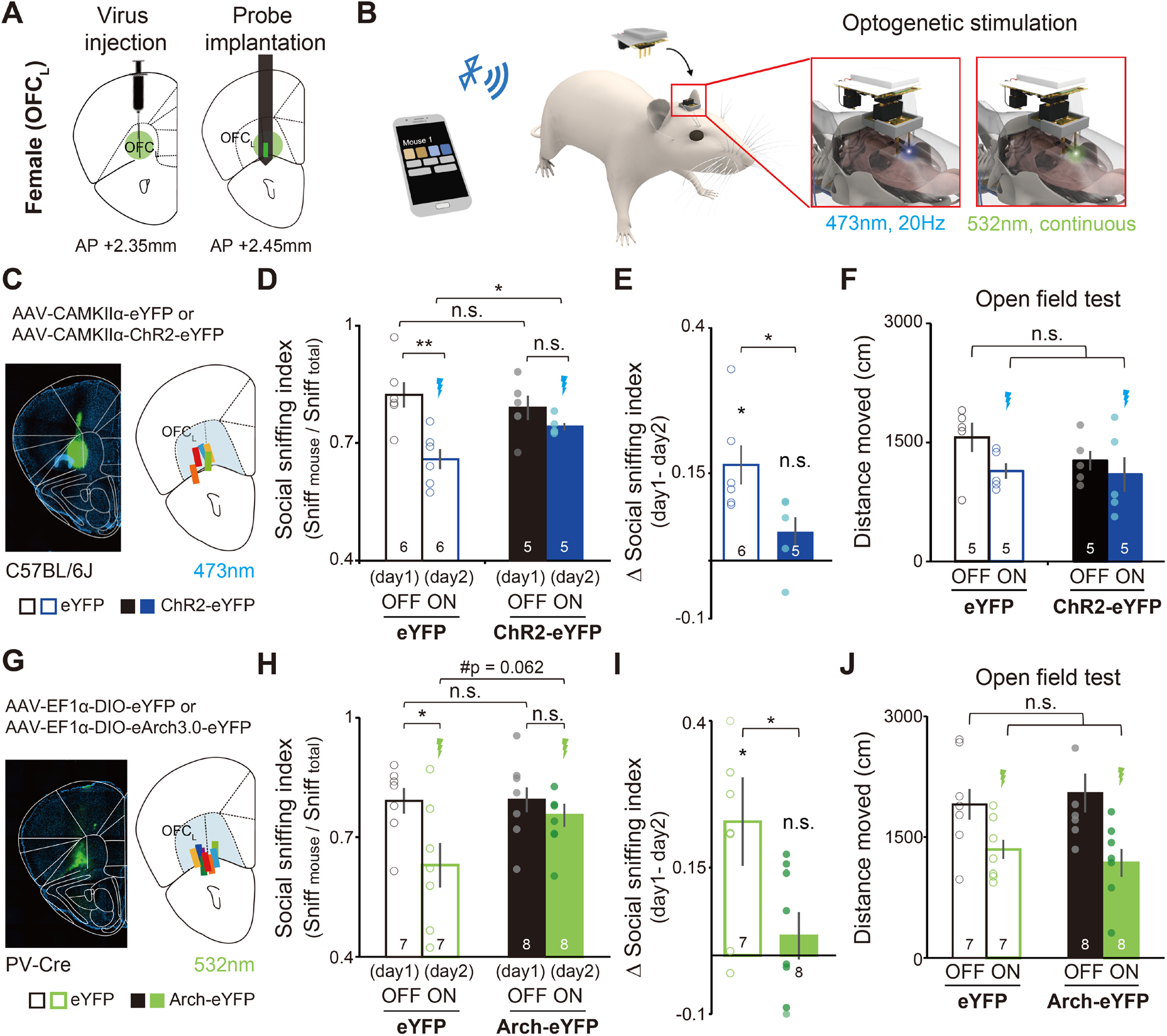
Optogenetic activation of excitatory neurons or inhibition of PV^+^ neurons in the OFC_L_ caused hypersociability in female mice. (A) Schematic illustration of virus injection and probe implantation for wireless optogenetic manipulation. The virus was injected into the OFC_L_ of a mouse. After a week of recovery, the wireless LED probe was implanted in the anterior part of OFC_L_ of a mouse. (B) Schematic illustration of wireless optogenetic manipulation. The smartphone application was used to turn on or off the light by Bluetooth Low Energy (BLE). For optogenetic stimulation, two types of LED light were used at different temporal modes: blue LED (473 nm) at a 20Hz light-pulse mode and green LED light (532nm) at a continuous light-on mode. (C) Optogenetic activation of excitatory neurons in the female OFC_L_. AAV-CAMKIIα-eYFP(eYFP, open bars) or AAV-CAMKIIα-ChR2-eYFP (ChR2-eYFP, filled bars) was injected into the OFC_L_ of PV-Cre female mice. Left, a representative brain image of the OFC_L_ of a WT female mouse injected with the AAV-CAMKIIα-ChR2-eYFP. Green, ChR2-eYFP; blue, DAPI. Right, a schematic illustration of a mouse brain section marked with each probe-implanted site in the OFC of ChR2-eYFP injected female mice. Blue shading, OFC; colored bars, probe-implanted sites confirmed by *post hoc* histology. (D) The social sniffing index of eYFP (open bars) and ChR-2eYFP (filled bars) mice was measured on two consecutive days. For Day1, the light was not delivered (OFF), and the social sniffing index was not different between the eYFP and ChR2-eYFP groups. Unpaired, Student’s t-test between groups, p = 0.516. For Day2, the light was delivered to the OFC_L_ (ON), and ChR2-eYFP group showed a higher social sniffing index than eYFP group. Unpaired, Student’s t-test between groups, *p = 0.030; Paired, Student’s t-test within a group, eYFP, **p = 0.007; ChR2-eYFP, p = 0.179. (E) Δ Social sniffing index indicates the difference in social sniffing index between day 1 (OFF) and day 2 (ON). Unpaired, Student’s t-test between groups, *p = 0.036; Wilcoxon signed rank test for each group, eYFP, *p = 0.036; ChR2-eYFP, p = 0.179. (F) Locomotion of mice (distance moved (cm) for 5 min) in the open field chamber was not affected by the optogenetic activation of the OFC_L_. Unpaired, Student’s t-test between groups, OFF, p = 0.267; ON, p = 0.874. Paired, Student’s t-test within a group, OFF vs ON, eYFP, p = 0.078; ChR2-eYFP, p = 0.602. Female, eYFP; *n* = 5 mice; ChR2, *n* = 5 mice. (G) Optogenetic inactivation of PV^+^ neurons in the female OFC_L_. AAV-EF1α-DIO-eYFP (eYFP) or AAV-EF1α-DIO-eArch3.0-eYFP (Arch-eYFP) was injected into the OFC_L_ of PV-Cre female mice. Left, a representative brain image of the OFC_L_ of a PV-Cre female mouse injected with the AAV-EFlα-DIO-eArch3.0-eYFP. Green, Arch-eYFP; blue, DAPI. Right, a schematic illustration of a mouse brain section marked with each probe-implanted site in the OFC of Arch-eYFP injected female mice. Blue shading, OFC; colored bars, probe-implanted sites confirmed by *post hoc* histology. (H) The social sniffing index of eYFP (open bars) and Arch-eYFP (filled bars) mice. For day 1, the light was not delivered (OFF), and the social sniffing index was not different between eYFP and Arch-eYFP groups. Unpaired, Student’s t-test between groups, day 1, p = 0.950; day 2, #p = 0.062. Paired, Student’s t-test within a group, eYFP, *p = 0.048; ChR2-eYFP, p = 0.480. (I) ΔSocial sniffing index of eYFP and Arch-eYFP. Unpaired, Student’s t-test between groups, *p = 0.046. Wilcoxon signed rank test for each group, eYFP, *p = 0.047; Arch-eYFP, p = 0.547. (J) Locomotion of mice (distance moved (cm) for 5 min) in the open field chamber was not affected by the optogenetic inactivation of the PV^+^ neurons in OFC_L_. Unpaired, Student’s t-test between groups, p = 0.068. Paired, Student’s t-test within a group, OFF vs ON, eYFP, **p = 0.007, Arch-eYFP, **p = 0.003. All data are presented as mean ± SEM. Dots indicate the individual data. The blue lightning sign indicates the delivery of light stimulation (20Hz, 473nm), and the green lightning sign indicates the delivery of light stimulation (continuous, 532nm). For (C-E), eYFP, *n* = 6; ChR2-eYFP, *n* = 5; female mice. For (G-J), eYFP, *n* = 7; Arch-eYFP, *n* = 8; female mice. **p < 0.01, *p < 0.05, #p < 0.1, n.s., not significant.

## Discussion

Our study found that adolescent social experience is critical for shaping sociability in female mice. Social isolation during late adolescence enhanced sociability in female mice while impairing the maturation of PV^+^ neurons and decreasing social sniffing-induced activity in the OFC_L_. The knock-down of PV in the OFC_L_ of female mice induced the same phenotype as the PWSI mice: enhanced sociability and decreased sniffing-induced activity in the OFC_L_. These effects were due to the imbalance between excitation and inhibition, as the PV knock-down caused hyper-excitation in the OFC_L_. We further found that optogenetic activation of excitatory neurons or suppression of PV^+^ inhibitory neurons in the OFC_L_ led to enhanced sociability in female mice. This finding supports the idea that the excitation-inhibition balance in the prefrontal cortex is critical for shaping social behaviors in mice (Yizhar et al., 2011). Our data also suggest that adolescent social isolation influences male and female mice differently, both in how the prefrontal cortex matures and social behavior matures.

### Importance of social experience during adolescence in shaping female social behavior

Social experience during adolescence is important for the maturation of the mammalian brain (Spear, 2000; Fuhrmann et al., 2015). In humans, social isolation during adolescence can increase the occurrence of neurological disorders and cause defects in brain function (Almeida et al., 2021). Similarly, depriving rodents of social experience during adolescence caused defects in the maturation of their brains and alterations in social and cognitive behaviors (Cooke et al., 2000; Schubert et al., 2009; Makinodan et al., 2012; Hinton et al., 2019; Potrebic et al., 2022). Furthermore, behavioral and physiological changes caused by adolescent social isolation were often sex-specific (Pisu et al., 2016; Bicks et al., 2020; Tan et al., 2021). In this study, we found that PWSI, particularly after puberty (from P50 to P80), impairs the maturation of inhibitory circuits in the OFC_L_ and causes maladaptation in social exploration behavior in female mice but not in male mice. Our study is unique compared to recent studies that examined the effects of social isolation during early adolescence (Bicks et al., 2020; Tan et al., 2021). Our findings emphasize the importance of later adolescence when females adjust and shape their social approaching behaviors with the maturation of PV^+^ neurons in the OFC_L_.

A recent study has shown that the 1dSI increased sociability in male mice (Matthews et al., 2016). We also found that male mice that had experienced 1dSI spent more time interacting with a novel mouse than did GH male mice (GH, 221.11 ± 17.93 s, 1dSI, 317.97 ± 31.46 s, p=0.023, student’s t-test). However, the 1dSI male mice also showed a tendency of increased interaction with an object in the other container (GH, 107.29± 9.88 s, 1dSI, 138.28 ± 23.36 s, p=0.256, student’s t-test). As a result, their social sniffing and social interaction indices, which indicate the ratio between social interaction and object interaction times, did not change in 1dSI male mice (Fig. **1C** and **1D**). Our data suggest that 1dSI may enhance object exploration behaviors in male mice. This possibility was not tested in the previous report, as they used an empty chamber without any object as opposed to the social chamber. Future studies are required to understand the effect of 1dSI in male mice fully. Regardless, our study focuses on the social isolation effect on female mice, and we found no effects of 1dSI on social and object interaction times in female mice. Our data demonstrate that long-term social isolation during late adolescence affects social behaviors more effectively than the 1dSI in females.

PWSI not only deprives sensory experience from social interaction but also enhances stress in social mammals like mice (Fone and Porkess, 2008). It has been shown that both sensory deprivation and stress can impair the maturation of PV^+^ neurons in the sensory cortex as well as in the hippocampus (Huang et al., 1999; Czeh et al., 2005; Hu et al., 2010). The OFC, which is classified as a secondary olfactory cortex, is one of the key areas involved in olfactory processing (Lipton et al., 1999) and sniffing behaviors (Kareken et al., 2004). Furthermore, the olfactory system is the most important way through which mice detect other mice (Radyushkin et al., 2009). The OFC has a strong connection with the olfactory system and the hippocampus (Wikenheiser and Schoenbaum, 2016). An fMRI study in humans showed that in response to odorant cues with emotional valence, only females showed increased activity in their OFC_L_ (Royet et al., 2003). Future studies are required to unravel which types of social olfactory cues and stress-related factors affect the maturation of PV^+^ neurons in the OFC_L_ and social behavior.

Our data suggest that socially evoked activity in the OFC_L_ is important for reducing social approaching behaviors when the animal becomes an adult. Excessive sociability is a symptom of patients with Williams-Beuren syndrome (Mimura et al., 2010; Sakurai et al., 2011; Segura-Puimedon et al., 2014; Stoppel and Anderson, 2017) or Angelman syndrome (Stoppel and Anderson, 2017; Simchi and Kaphzan, 2021), rare neurodevelopmental genetic disorders. Interestingly, these patients show abnormal activity patterns in the OFC. Hypersociability is also found in some mouse models of autism (Yoo et al., 2019). Our data demonstrate that PWSI induced similar behavioral abnormalities in female mice, suggesting that the normal maturation and activity of the OFC_L_, at least in females, is important for maintaining an adequate level of sociability in such disorders.

### PV expression in the OFC_L_ is important for female social behavior

Recent studies found important roles of PV expression in the PFC in shaping social and cognitive behaviors. For example, several mouse models of autism that show impaired sociability have shown reduced PV expression in the mPFC (Filice et al., 2016; Kobayashi et al., 2018). Furthermore, maternal separation before weaning (Goodwill et al., 2018) or social isolation during early adolescence (Hinton et al., 2019) reduced PV expression in the OFC and impaired flexibility in behavioral decisions in rodents. Whole-brain knock-out of PV caused the mice to show autism-like behavior (Wohr et al., 2015), but it is still unclear whether the reduced PV expression in sub-regions of the PFC caused such behavioral deficits. Our study newly applied the CRISPR/Cas9 system to reduce PV expression locally in the OFC_L_. Using this method, we confirmed that PV expression in the OFC_L_ during adolescence is critical for shaping social behaviors in female mice (Fig. **5**). This technique can be widely used for studying area-specific PV protein function in various brain areas, although it lacks the reversibility of the loss-of-function by permanent gene deletion *in vivo*.

PV is a calcium-binding protein that plays a key role in regulating the physiological properties of PV^+^ neurons. In our study, knocking down PV decreased the excitability of the PV^+^ inhibitory neurons, which in turn decreased synaptic inhibition on nearby excitatory neurons (Fig. **6H-6M**). The relative reduction in inhibition compared to excitation in the OFC_L_ potentially disrupted overall social sniffing-induced activity in the OFC_L_ (Fig. **6F** and **6G**) and hypersociability (Fig. **5H**). We further proved this idea by the optogenetic activation of excitatory neurons in the OFC_L_ that led to hypersociability (Fig. **7**). Moreover, a reduced number of PV^+^ neurons can also disrupt the network property of the OFC_L_, similar to the previous report showing that the loss of PV^+^ neurons correlated with a decrease in the network oscillation in the PFC (Lodge et al., 2009). Maturation of network activity during adolescence may be critical for shaping social sniffing-induced activity in the OFC_L_.

In a study of virgin female mice, PV and PNN expression in the auditory cortex was reported to be correlated with learning to retrieve pups (Krishnan et al., 2017); this activity, which occurs in the left auditory cortex, is important for shaping maternal behaviors in female mice (Marlin et al., 2015). Although several studies have shown that the expression of PNN is important for the maturation of PV^+^ neurons (Gogolla et al., 2009), we found expression of the PV but not the PNN is important for the maturation of the female OFC_L_ during late adolescence. Our data suggest that transcription or translation of PV and PNN may be separately regulated, and adolescent social isolation specifically disrupted PV expression only in a subset of neurons in the OFC_L_. Our data further support the idea that plasticity in PV^+^ neurons in the OFC_L_ is important for shaping female sociability and is also characterized by hemispheric asymmetry. Both genetic and epigenetic mechanisms have been identified as critical for regulating normal social behavior (Hensch, 2005), and thus social behaviors can be easily disrupted in a wide range of brain psychiatric disorders (Blakemore, 2008). Our study further provides insight into how disruption of experience-dependent gene expression in the PFC inhibitory neurons, such as reduced PV expression in the OFC_L_, may underlie abnormal hypersociability that characterizes certain psychiatric disorders.

## Acknowledgments

This work was supported by grants from the National Research Foundation of Korea (2021R1A2C3012159, 2021R1A4A2001803, to S.-H.L. and 2021R1A2C4001483, to J.-W.J.) and the KAIST Global Singularity Program for 2020 (to S.-H.L.). This work was also supported by the Rural Development Administration (PJ016403, to S.K.) and by the Institute for Basic Science (IBS-002-D1 to E.K. and IBS-002-D3 to S.-H.L.). We thank all members of the Lee laboratory for their helpful discussion, Dr. Soo Young Kim for discussion on the PWSI, and Emily Wheeler, Boston, for editorial assistance.

## Contact for reagent and resource sharing

Further information about and requests for reagents may be directed to and will be fulfilled by the corresponding author, Dr. Seung-Hee Lee.

## Author contributions

YSJ and DJ performed most of the experiments and contributed equally to this work; HK and EK performed and evaluated patch-clamp recording experiments; JHK helped to perform *in vivo* recordings and measure sniffing behavior; CYK and JWJ designed and fabricated the wireless-controllable optoelectronic neural probes; YO and SGK designed and cloned the AAV-sgPV vectors; YHL and CHK cloned the AAV-tdTomato vectors; SHL designed and supervised the study. YSJ, DJ, SHL wrote the manuscript with contributions from all the authors.

